# Divergent associations of slow-wave sleep vs. REM sleep with plasma amyloid-beta

**DOI:** 10.1101/2023.06.02.543111

**Authors:** Yevgenia Rosenblum, Mariana Pereira, Oliver Stange, Frederik D. Weber, Leonore Bovy, Sofia Tzioridou, Elisa Lancini, David A. Neville, Nadja Klein, Timo de Wolff, Mandy Stritzke, Iris Kersten, Manfred Uhr, Jurgen A.H.R. Claassen, Axel Steiger, Marcel M. Verbeek, Martin Dresler

## Abstract

**Background:** Recent evidence shows that during slow-wave sleep (SWS), the brain is cleared from potentially toxic metabolites, such as the amyloid-beta protein. Poor sleep or elevated cortisol levels can worsen amyloid-beta clearance, potentially leading to the formation of amyloid plaques, a neuropathological hallmark of Alzheimer’s disease. Here, we explore how nocturnal neural and endocrine activity affects amyloid-beta fluctuations in the peripheral blood as a reflection of cerebral clearance.

**Methods:** Simultaneous polysomnography and all-night blood sampling were acquired in 60 healthy volunteers aged 20–68 years old. Nocturnal plasma concentrations of two amyloid-beta species (amyloid-beta-40 and amyloid-beta-42), cortisol, and growth hormone were assessed every 20 minutes from 23:00–7:00. Amyloid-beta fluctuations were modeled with sleep stages, (non)-oscillatory power, and hormones as predictors while controlling for age and multiple comparisons. Time lags between the predictors and amyloid-beta ranged from 20 to 120min.

**Findings:** The amyloid-beta-40 and amyloid-beta-42 levels correlated positively with growth hormone concentrations, SWS proportion, slow-wave (0.3–4Hz) oscillatory and high-band (30–48Hz) non-oscillatory power, but negatively with cortisol concentrations and rapid eye movement sleep (REM) proportion measured 40–100min before (all t-values>|3|, p-values<0.003). Older participants showed higher amyloid-beta-40 levels.

**Interpretation:** Slow-wave oscillations are associated with higher plasma amyloid-beta levels, reflecting their contribution to cerebral amyloid-beta clearance across the blood-brain barrier. REM sleep is related to decreased amyloid-beta plasma levels; however, this link may reflect passive aftereffects of SWS and not REM’s effects per se. Strong associations between cortisol, growth hormone, and amyloid-beta presumably reflect the sleep-regulating role of the corresponding releasing hormones. A positive association between age and amyloid-beta-40 may indicate that peripheral clearance becomes less efficient with age. Our study provides important insights into the specificity of different sleep features’ effects on brain clearance and suggests that cortisol nocturnal fluctuations may serve as a new marker of clearance efficiency.

## Introduction

In recent years, a metabolic homeostatic function of sleep has been put forward. In rodents, an increase in the interstitial space surrounding tissue cells has been observed during sleep, resulting in an increased clearance of potentially neurotoxic metabolic products, particularly, the amyloid-beta (Aβ) protein^1^. In humans, experimental sleep deprivation or interference with slow-wave sleep (SWS) inhibits the overnight reduction in Aβ cerebrospinal fluid (CSF), thereby increasing CSF Aβ concentrations^2,3,4^ while self-reported sleep quality is linked to Aβ deposition in cognitively healthy older adults^5^.

The important role of sleep in Aβ clearance is further supported by the facts that in Alzheimer’s disease (AD), impaired sleep and pathological accumulation of cortical Aβ are reported long before (sometimes up to 20 years) the onset of clinical symptoms^6^. Moreover, it has been suggested that impaired sleep can lead to Aβ accumulation which in turn can drive the progression of the disease^6^.

Other lines of evidence report that Aβ clearance can be affected by abnormally increased cortisol concentrations, another hallmark of AD^7^. High levels of circulating cortisol are usually associated with sustained stress. Within a normal circadian cycle, cortisol levels increase during rapid eye movement (REM) and decrease during SWS, which is further correlated with a surge of growth hormone (GH)^8^. These associations suggest common regulators of neural and endocrine activity during sleep and possibly of the endocrine and metabolic functions of sleep.

This study explores how sleep features, cortisol, and GH affect subsequent Aβ plasma levels as a reflection of cerebral clearance. To achieve this aim, we acquire concurrent all-night plasma sampling and polysomnography in a large sample of healthy participants. Based on the literature, we hypothesize that SWS and GH peak (which temporally coincides with SWS) would be linked to an increase in the subsequent Aβ plasma levels. Given that REM sleep is accompanied by wake-like neural activity^10^, we hypothesize that the REM episodes and the cortisol peak (which temporally coincides with REM) are accompanied by cerebral waste accumulation. However, we have no prior hypothesis on the direction of the correlation (if any) between REM sleep and plasma Aβ since the existence of Aβ clearance during REM sleep has not been reported in the literature so far.

## Methods

### Participants

We analyzed plasma samples and polysomnographic recordings from previous unpublished endocrinological studies conducted at the Max Planck Institute of Psychiatry, using only nights with no pharmacological or endocrine intervention. The sample consisted of 60 healthy participants (32 females) aged 39.8±16.0 years (range 20–68). All studies and data re-analyses were approved by the Ethics committee of the University of Munich. All participants gave written informed consent.

### Plasma measurement

In the experimental night in the sleep laboratory, 4 ml blood were drawn every 30 min (20:00– 22:00 h) or 20 min (22:00–07:00 h), respectively, from the adjacent room, using an intravenous cannula and a tube extension. Free GH and cortisol concentrations were measured by radioimmunoassay. Aβ1-40 and Aβ1-42 were analyzed with commercially available Euroimmun ELISAs (plasma protocol^9^) according to the manufacturer’s instructions by an expert blind to the subject characteristics and sleep data. Only the samples drawn from 23:00 to 7:00 were analyzed (Fig.S1). All samples from one subject were measured on the same ELISA plate in the same run.

### Polysomnography

The experimental night was preceded by an adaptation night in the sleep laboratory. Polysomnography was recorded from 23:00 to 07:00, stored, and analyzed with a digital recorder (Comlab 32 Digital Sleep Lab, Brainlab V 3.3 Software, Schwarzer GmbH, Munich, Germany) from F3, F4, C3, C4, P3, P4, O1, and O2 leads, electrooculogram, and mental/submental electromyogram, with a sampling rate of 250 Hz (filtered from 0.3 to 70 Hz). Sleep data was scored according to the AASM standards by experts not involved in the study. Epochs with artifacts were detected by visual inspection and rejected from further analysis.

### Spectral power

Total spectral power was calculated for each 30s epoch for each channel and then differentiated to its aperiodic (i.e., fractal, 1/f, scale-free) and oscillatory components using the Irregularly Resampled Auto-Spectral Analysis^10^. To implement the algorithm, we used the *ft_freqanalysis* function of the Fieldtrip toolbox^11^ as described elsewhere^12,13^. The function was called twice, with the *cfg*.*output=‘fractal’* and *cfg*.*output=‘original’* for the total power and its aperiodic component, respectively. The aperiodic power component was transformed to log-log coordinates by standard least-squares regression. To estimate the power-law exponent β (the rate of the spectral decay), we calculated the slope of the aperiodic component as a marker of excitation-to-inhibition ratio^14^. The oscillatory component was calculated by subtracting the aperiodic component from the total power. All values higher than 3 standard deviations above the mean were automatically replaced by NaN values.

### Sleep outcome measures

Sleep features were characterized with six different variables; specifically, the proportion of N1, N2, SWS, and REM sleep, oscillatory power component in the 0.3-4Hz band (slow-wave activity, SWA) as an objective EEG marker of SWS, and the slope of the aperiodic power component in the 30–48Hz band as an objective marker of REM sleep^15^ (Fig.S4). The analysis was limited to 48Hz due to recording line noise (50Hz in Europe). All variables were averaged over each 20 min of sleep. Spectral power variables were averaged over F3 and F4 electrodes, as this location is associated with the highest SWA.

As an exploratory sub-analysis, we also calculated the oscillatory power component in the theta (4–8Hz), alpha (8–11Hz), and beta (15–30Hz) frequency bands, as well as the slope of the aperiodic power component in the 0.3–35Hz band and averaged them over the frontal electrodes.

In Supplementary Materials, we report the effects of spectral power over additional topographical areas (S1), atonia (S2), and autonomic functioning (S3) on Aβ clearance.

### Statistical analysis

To assess whether sleep features or hormones (predictors) can explain changes in the subsequent Aβ1-40 or Aβ1-42 plasma levels (dependent variables) we used univariable mixed-effects models corrected for participant’s age and subject using the R package *mgcv*.

Specifically, each model includes an age effect, a participant-specific random intercept (to control for individual-specific heterogeneities) and one of the sleep features or hormones. The code is provided in Supplementary Material S4.

Since an *a priori* hypothesis regarding the time lag between the predictors and dependent variable was not possible due to the lack of relevant findings in the literature, for each sleep feature/hormone, we ran six models corresponding to the time lags between the predictor and Aβ ranging from -120 to -20 min with a 20-min step (Fig.S1). To control for multiple comparisons, we used Benjamini-Hochberg’s adjustment with a false discovery rate set at 0.05. Due to the semi-exploratory nature of this study, all corrections were done per a given sleep feature/hormone (six different tests) with the α-level set in the 0.008-0.050 range.

To assess model quality, we calculated the R^2^ and Akaike information criterion (AIC). Given that the number of observations ranged from 19 per subject for the models using the 120-min lag to 24 per subject for the models using the 20-min lag, we excluded the last 1–5 observations from the longer datasets to end up with an equal number of observations (∼19 per subject) in each model. We note that results are not directly comparable across time lags as they are not based on the exact same set of data points. Nevertheless, assuming that the entire sample is representative, our models can serve as initial descriptive visualizations able to hint which time lag should be chosen for the next-level analysis.

## Results

The demographic, sleep, and plasma characteristics of the participants are reported in Table 1. Averaged nocturnal Aβ and hormone plasma concentrations are shown in Fig.1A. The models that used the –80-min lag between the predictors and Aβ showed the lowest AICs (i.e., the best models). They are presented in Fig.2 and Table 2. The slope estimates of all linear models are presented in Fig.3.

**Table 1:**
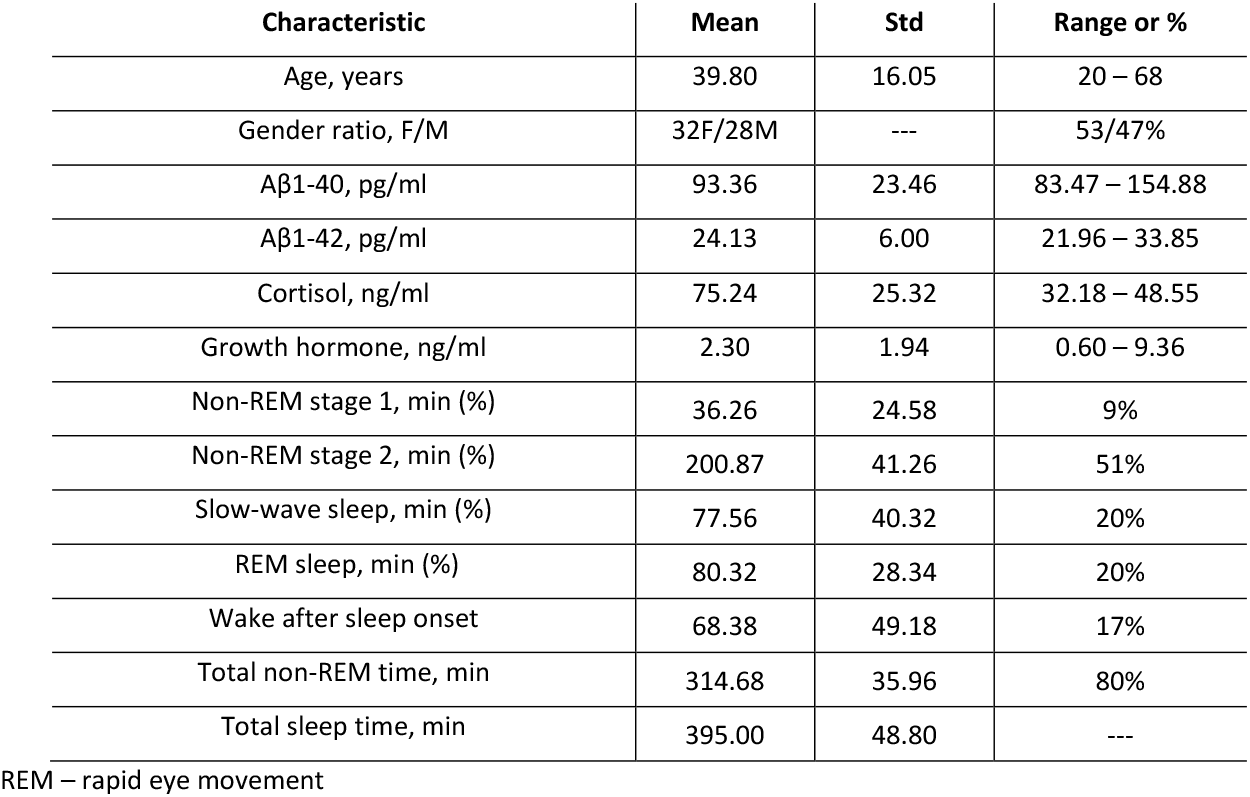
Demographic, plasma, and sleep characteristics of the participants.

**Figure 1.**
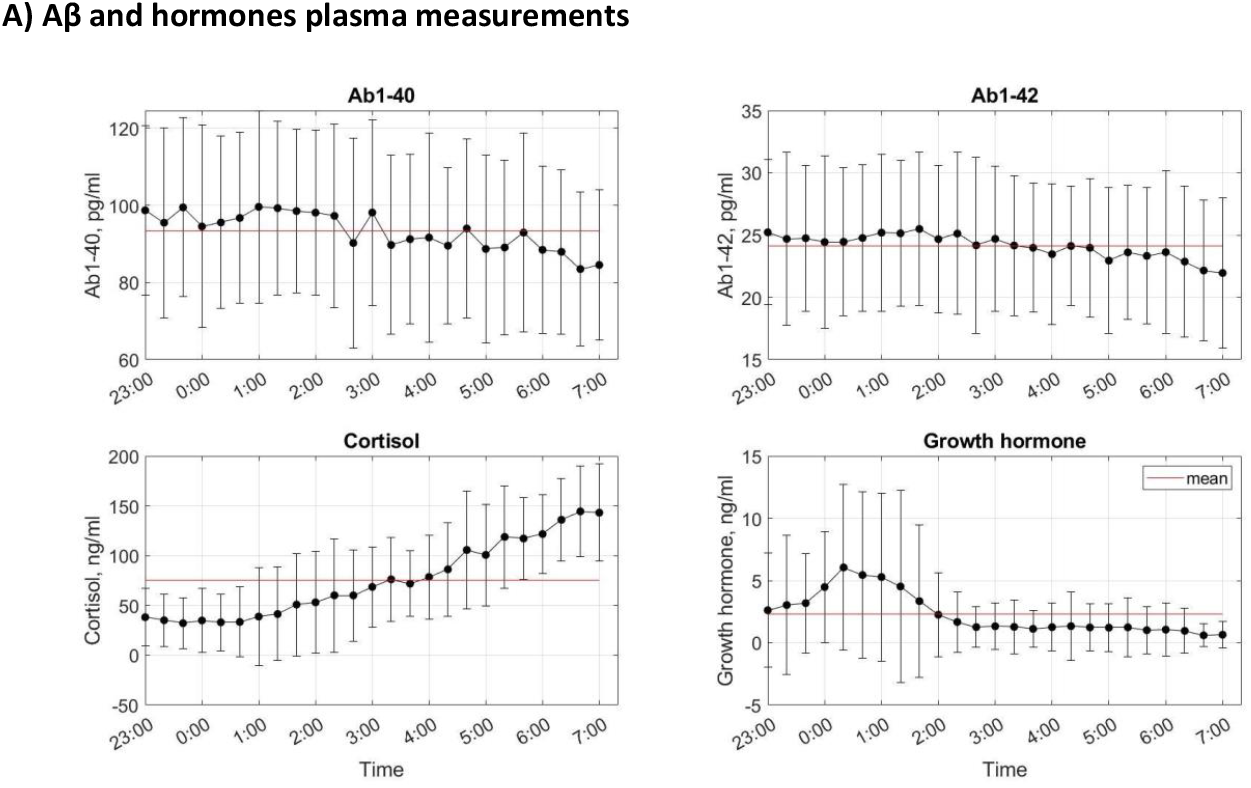
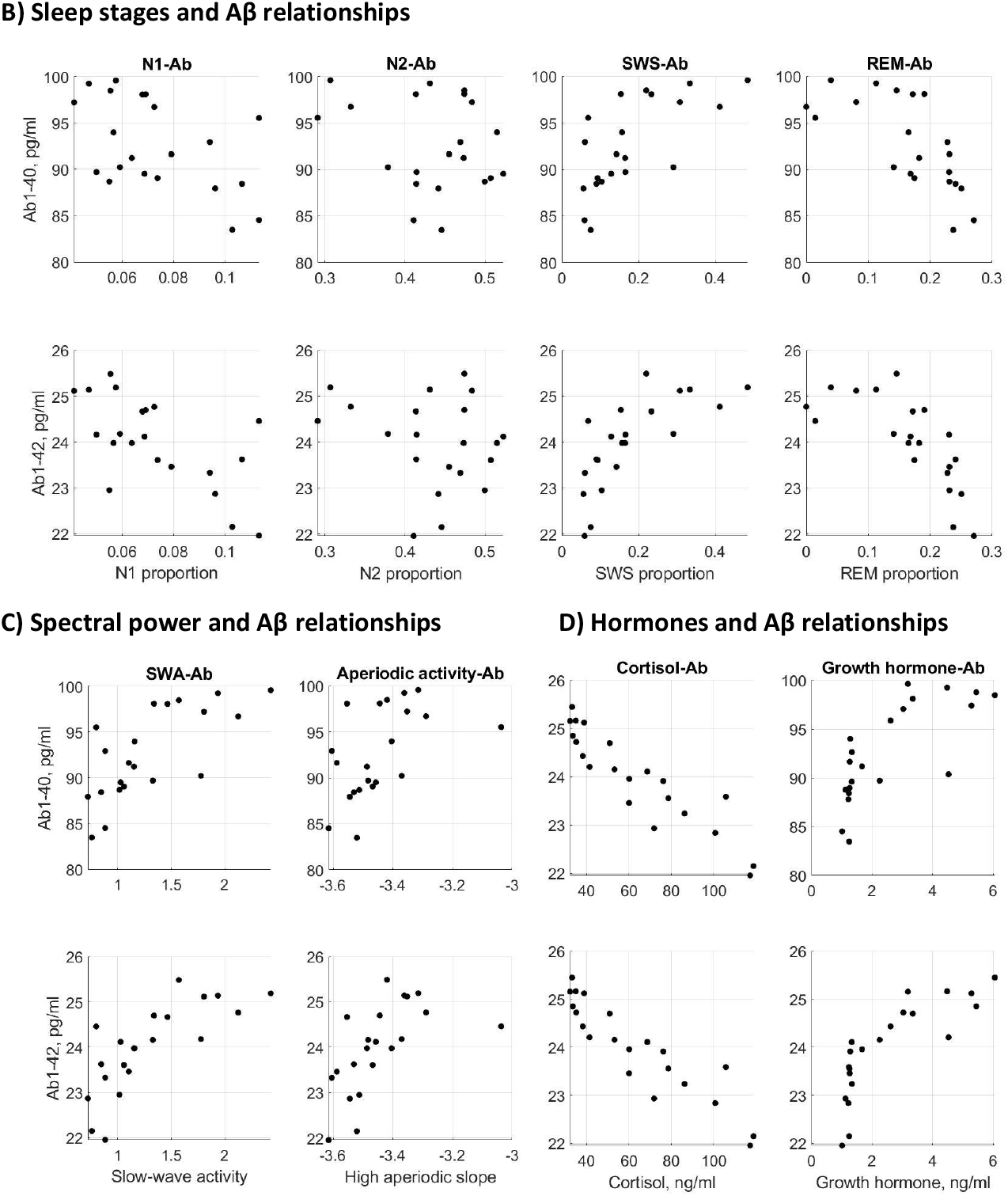
Aβ, hormones and sleep features. **(A)** Averaged Aβ, cortisol and growth hormone plasma concentrations with SD measured each 20 min from 23:00 till 7:00. Error bars denote standard deviations between the individuals. On average, plasma Aβ levels decreased by 13– 15% across the night. The cortisol profile increased towards morning, whereas growth hormone showed a prominent peak during the first part of the night and then decreased. **(B-D)** The relationships between the group-level averaged sleep stages (B), spectral power (C), and hormones (D) (used as predictors in the statistical models shown in Fig.2–3) and Aβ measured 80 min thereafter. SWS – slow-wave sleep, SWA – slow-wave activity, REM – rapid eye movement.

**Table 2:**
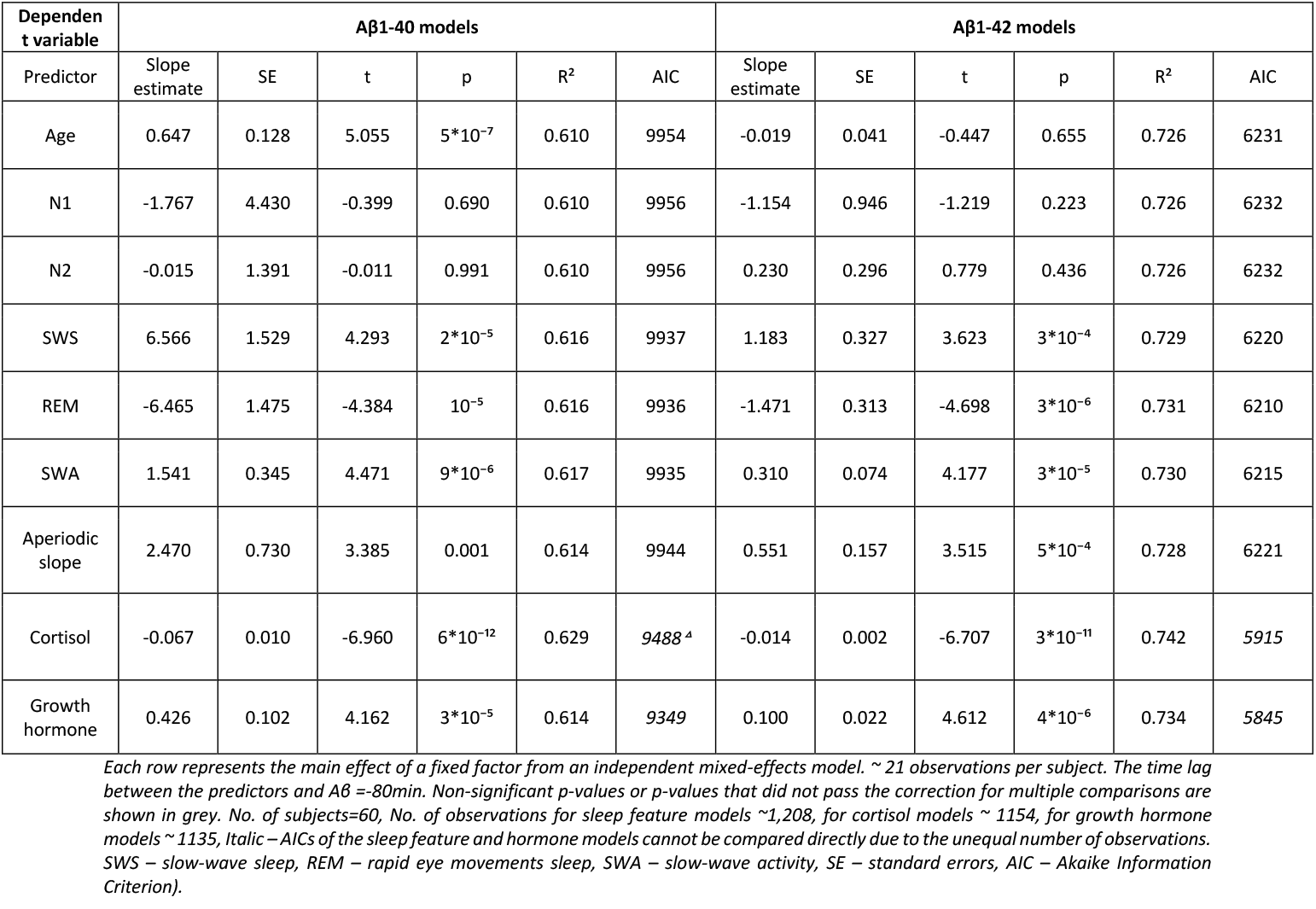
Linear mixed-effects models.

| Dependent variable | Aβ1-40 models | | | | | | Aβ1-42 models | | | | | |
| --- | --- | --- | --- | --- | --- | --- | --- | --- | --- | --- | --- | --- |
| Predictor | Slope estimate | SE | t | p | R<sup>2</sup> | AIC | Slope estimate | SE | t | p | R<sup>2</sup> | AIC |
| Age | 0.647 | 0.128 | 5.055 | 5*10<sup>-7</sup> | 0.610 | 9954 | -0.019 | 0.041 | -0.447 | 0.655 | 0.726 | 6231 |
| N1 | -1.767 | 4.430 | -0.399 | 0.690 | 0.610 | 9956 | -1.154 | 0.946 | -1.219 | 0.223 | 0.726 | 6232 |
| N2 | -0.015 | 1.391 | -0.011 | 0.991 | 0.610 | 9956 | 0.230 | 0.296 | 0.779 | 0.436 | 0.726 | 6232 |
| SWS | 6.566 | 1.529 | 4.293 | 2*10<sup>-5</sup> | 0.616 | 9937 | 1.183 | 0.327 | 3.623 | 3*10<sup>-4</sup> | 0.729 | 6220 |
| REM | -6.465 | 1.475 | -4.384 | 10<sup>-5</sup> | 0.616 | 9936 | -1.471 | 0.313 | -4.698 | 3*10<sup>-6</sup> | 0.731 | 6210 |
| SWA | 1.541 | 0.345 | 4.471 | 9*10<sup>-6</sup> | 0.617 | 9935 | 0.310 | 0.074 | 4.177 | 3*10<sup>-5</sup> | 0.730 | 6215 |
| Aperiodic slope | 2.470 | 0.730 | 3.385 | 0.001 | 0.614 | 9944 | 0.551 | 0.157 | 3.515 | 5*10<sup>-4</sup> | 0.728 | 6221 |
| Cortisol | -0.067 | 0.010 | -6.960 | 6*10<sup>-12</sup> | 0.629 | 9488<sup>a</sup> | -0.014 | 0.002 | -6.707 | 3*10<sup>-11</sup> | 0.742 | 5915 |
| Growth hormone | 0.426 | 0.102 | 4.162 | 3*10<sup>-5</sup> | 0.614 | 9349 | 0.100 | 0.022 | 4.612 | 4*10<sup>-6</sup> | 0.734 | 5845 |

Each row represents the main effect of a fixed factor from an independent mixed-effects model. ~ 21 observations per subject. The time lag between the predictors and Aβ = -80min. Non-significant p-values or p-values that did not pass the correction for multiple comparisons are shown in grey. No. of subjects=60, No. of observations for sleep feature models ~1,208, for cortisol models ~ 1154, for growth hormone models ~ 1135, *Italic – AICs of the sleep feature and hormone models cannot be compared directly due to the unequal number of observations.* SWS – slow-wave sleep, REM – rapid eye movements sleep, SWA – slow-wave activity, SE – standard errors, AIC – Akaike Information Criterion).

### Plasma measurements

Across the night, the Aβ1-40 levels decreased by 15% from 99.1±22.1 pg/ml at 23:00 to 84.5±19.5 pg/ml at 7:00. The Aβ1-42 levels decreased by 13% from 25.2±5.8 pg/ml at 23:00 to 21.9±6.0 pg/ml at 7:00. The cortisol concentrations showed a 375% increase from 23:00 to 7:00. The GH levels showed a prominent peak during the first part of the night (a 232% increase compared to the baseline measurement at 23:00) and then decreased. These findings replicated the existing literature.

### Age

We found that after the correction for multiple comparisons (6 tests), only the pre-sleep (measured at 23:00) and mean, but not post-sleep (measured at 7:00) levels of Aβ1-40 positively correlated with the participants’ age. The post-sleep Aβ1-42 levels negatively correlated with the participants’ age; however, this correlation did not pass the correction (Fig. S3). The mixed-effects models confirmed that participants’ age predicted higher plasma levels of Aβ1-40 but not Aβ1-42 (Table 2, Fig.2).

**Figure 2.**
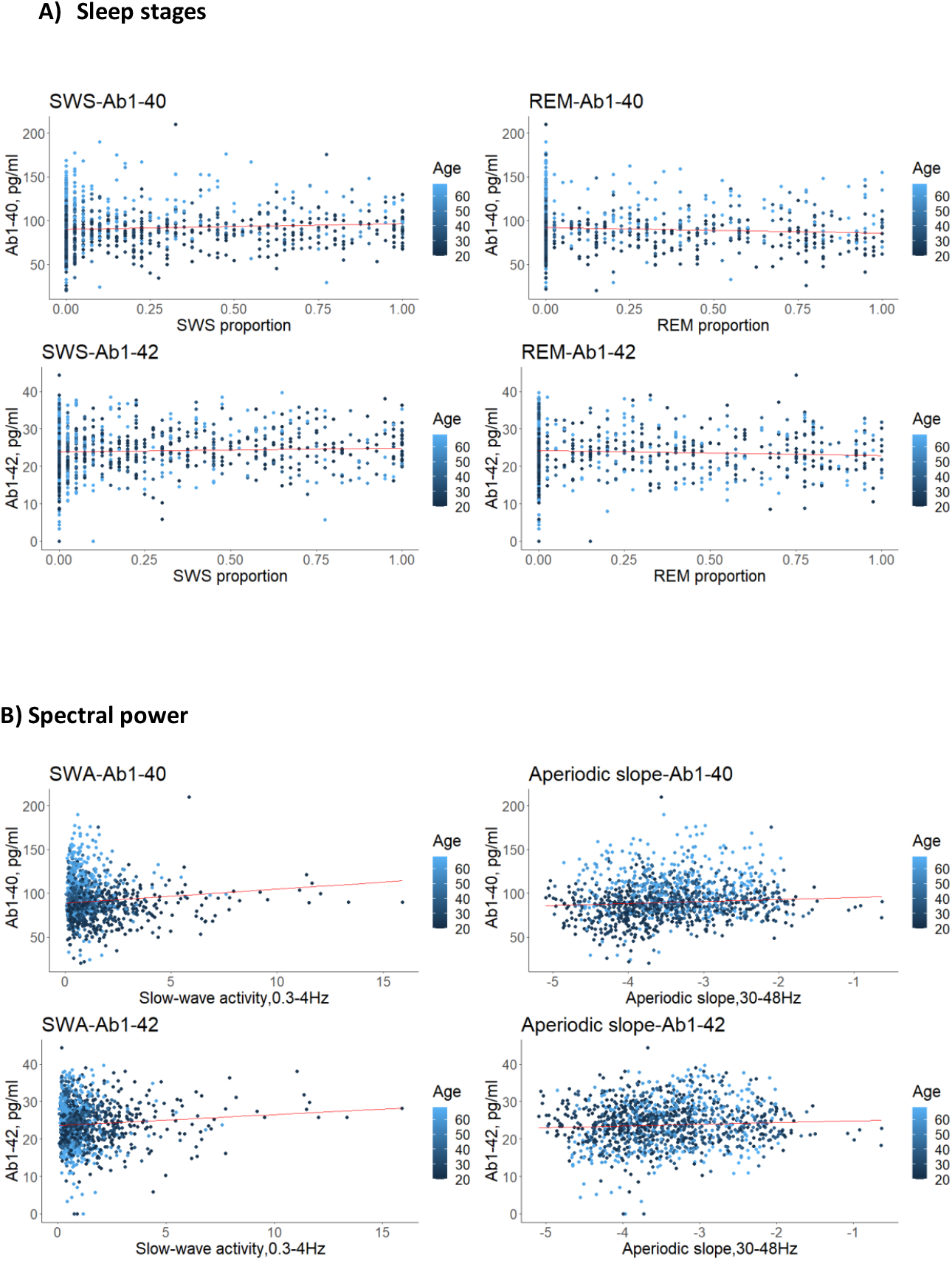
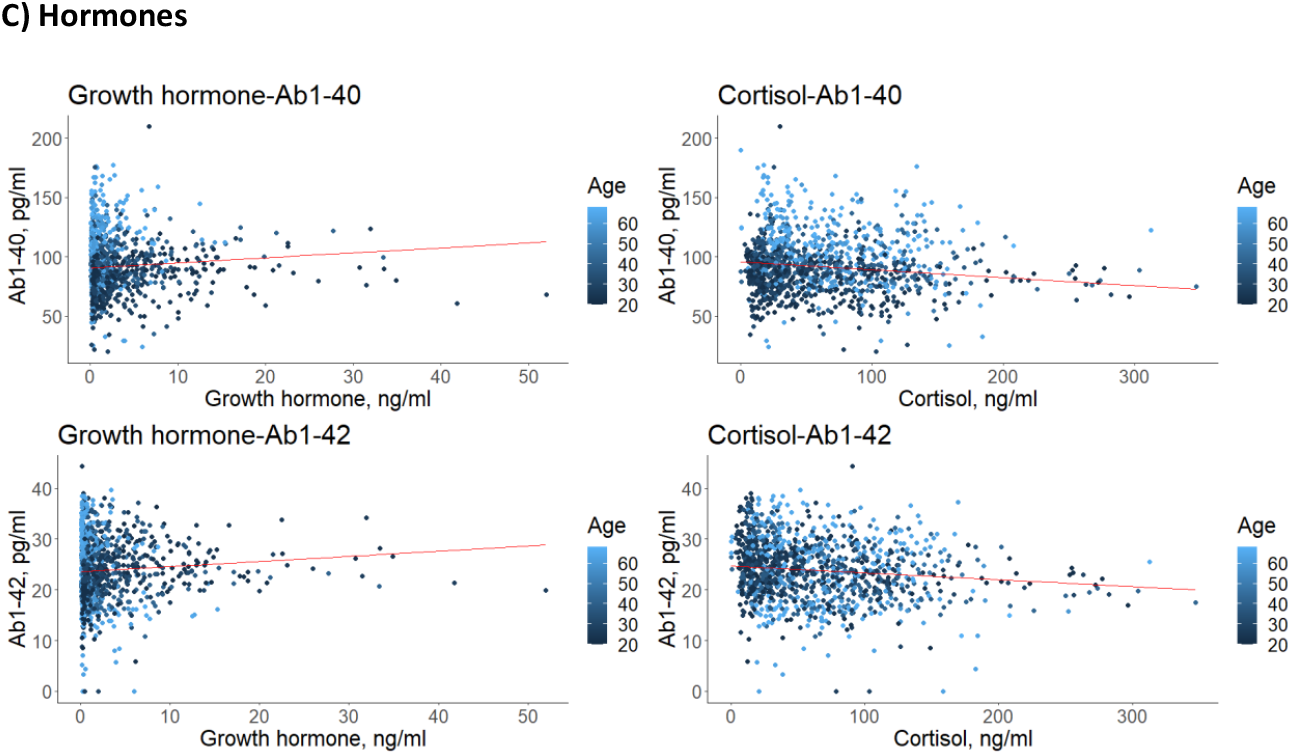
Mixed-effects models. **(A)** The SWS proportion, REM sleep proportion, **(B)** frontal SWA (0.3-4Hz), the slopes of the frontal high-band aperiodic power (30-48Hz), **(C)** cortisol or growth hormone levels were entered into (separate) linear (red lines) mixed-effects models as fixed factors together with the “age” as an additional fixed factor and the “participant” as a random factor to predict the plasma levels of Aβ1-40 and Aβ1-42 measured 80 min thereafter. Within an 80-minutes delay the Aβ1-40 and Aβ1-42 plasma levels correlated positively with growth hormone concentrations, SWS proportion, frontal SWA, aperiodic high-band slope, but negatively with cortisol concentrations and REM sleep proportion. Younger age (darker dots) predicts lower plasma levels of Aβ1-40 (as reflected by higher concentrations of the darker dots in the lower part of the graphs), whereas older age (lighter dots) predicts higher plasma levels of Aβ1-40 (as reflected by higher concentrations of the lighter dots in the higher part of the graphs). SWS – slow-wave sleep, SWA – slow-wave activity, REM – rapid eye movement.

**Figure 3.**
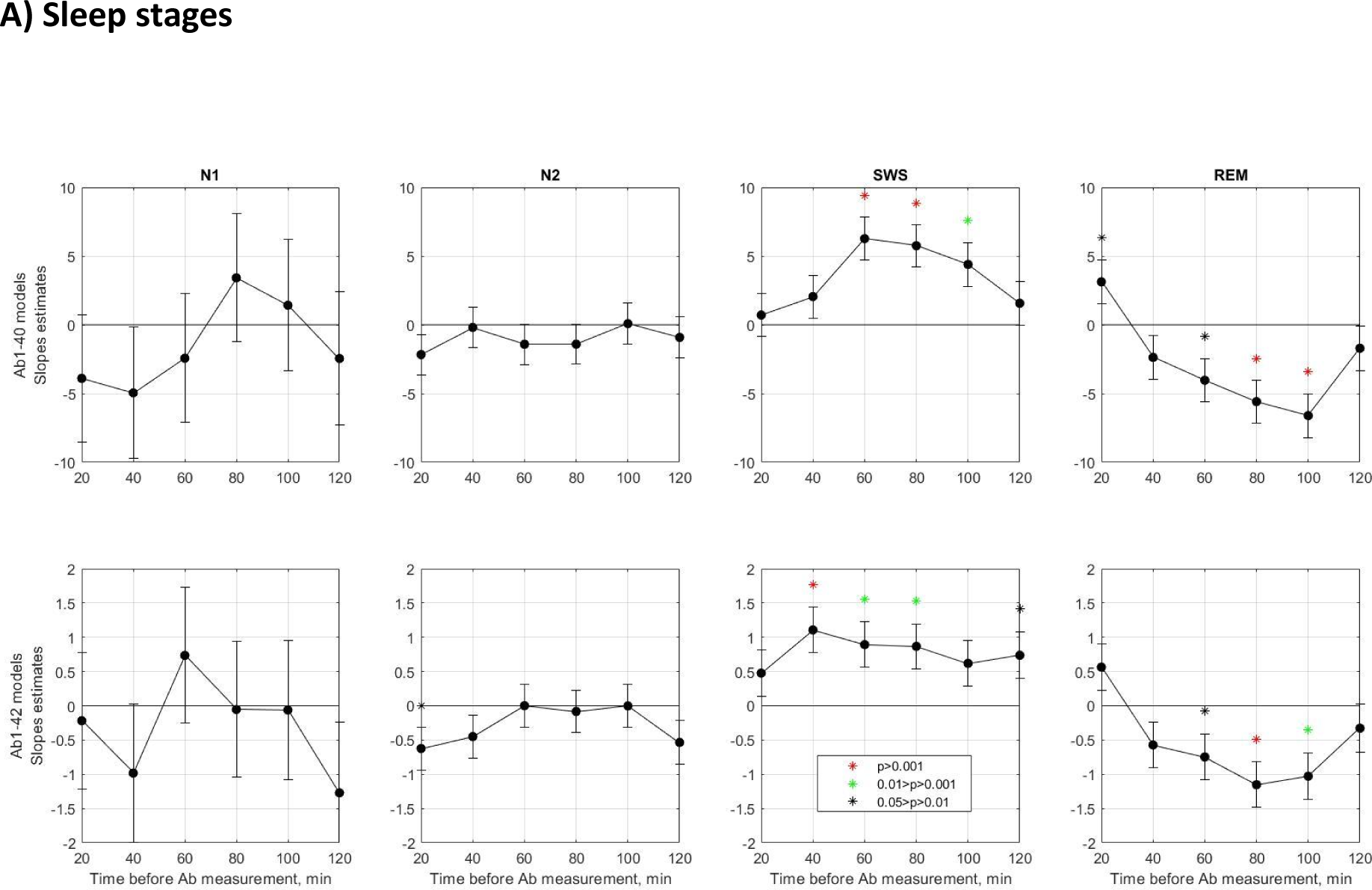
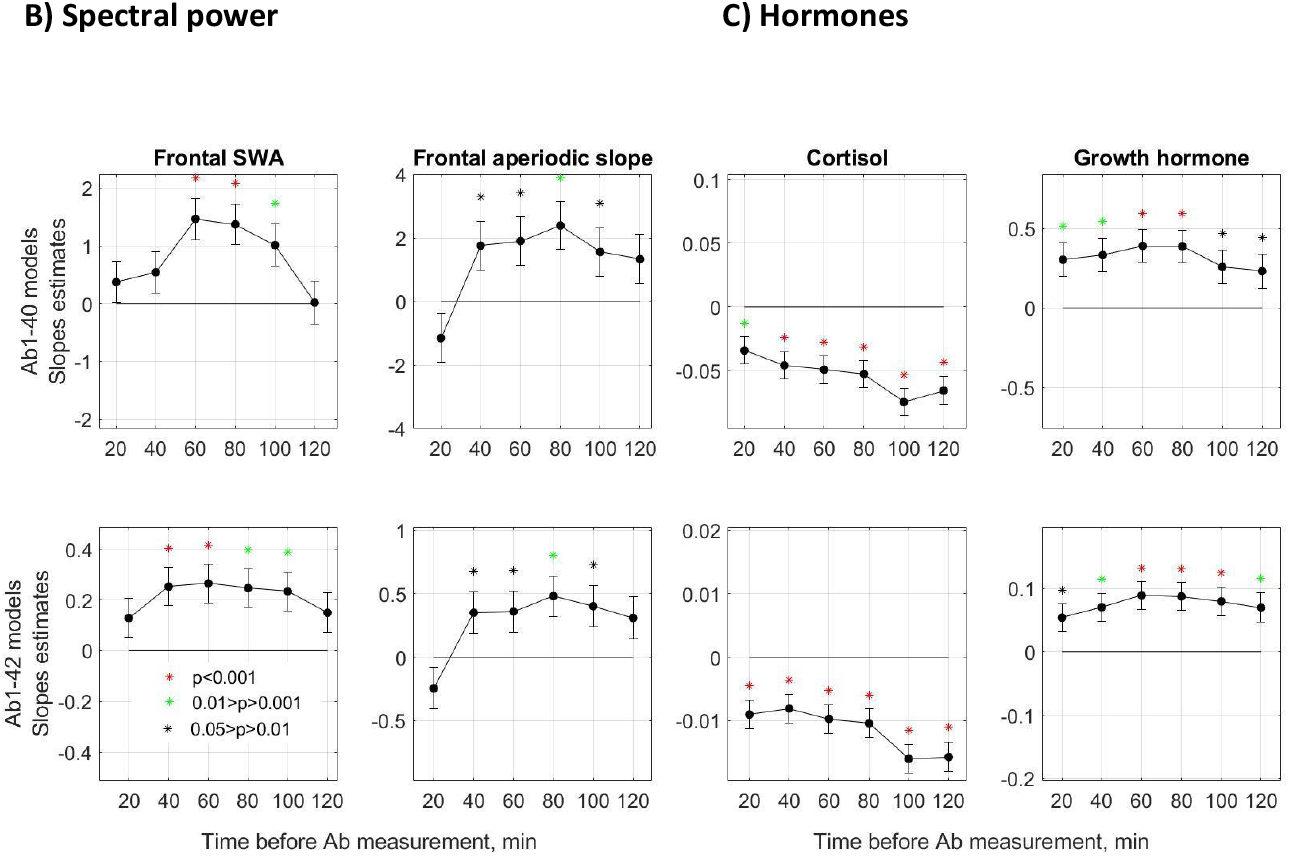
Slope estimates of the Aβ predictors. **(A)** The proportion of N1, N2, SWS, or REM sleep, **(B)** frontal spectral power, or **(C)** cortisol or growth hormone levels were entered as predictors of Aβ1-40 and Aβ1-42 into 6 different linear mixed-effects models using time lags ranging from -20 to -120 min between the predictors and Aβ. The slope estimate of the fixed effect of each predictor and its SE are presented. The asterisks mark the slope estimates for which the null hypothesis (stating that a predictor does not significantly affect the response (Aβ)) should be rejected. The models that passed the correction for multiple comparisons are listed in the Main Text. The SWS proportion, SWA, GH, and high aperiodic slope predict an increase in the subsequent Aβ levels (significantly positive slopes), the REM sleep proportion and cortisol predict a decrease (significantly negative slopes) in the subsequent Aβ levels, whereas N1 and N2 have no effect (close to zero slopes) on the subsequent Aβ levels. SWS – slow-wave sleep, SWA – slow-wave activity, REM – rapid eye movement sleep. Red asterisks correspond to p<0.001, green – to 0.01>p>0.001, and black – to 0.05>p>0.01.

### Sleep stages

SWS proportion predicted higher Aβ1-40 measured 40–100 min thereafter and Aβ1-42 measured 20–80 and 100–120 min thereafter. Rapid eye movement (REM) sleep proportion measured at 40-100 min prior to a blood measurement predicted lower subsequent levels of Aβ1-40 and Aβ1-42. N1 and N2 had no effect on Aβ (Fig.3).

### Spectral power

SWA (an EEG marker of SWS) predicted higher Aβ1-40 measured 40–100 min afterward and Aβ1-42 measured 20–100 min afterward (Fig.3). Oscillatory spectral power in the theta, alpha, and beta bands had no effect on Aβ. The high band (30–48Hz) aperiodic slopes (an EEG marker of REM sleep) predicted higher Aβ1-40 measured 20–80 min later and Aβ1-42 measured 20–100 min afterward (Fig.3). The low-band (0.3–35Hz) aperiodic slopes had no effect on Aβ.

### Hormones

Cortisol correlated negatively while GH correlated positively with Aβ1-40 and Aβ1-42 measured 0–120 min thereafter (Fig.3).

## Discussion

The present study investigated how nocturnal neural and endocrine activity affects Aβ clearance using a large sample of healthy individuals aged 20–68 years old. The results show that the Aβ1-40 and Aβ1-42 plasma levels correlate positively with growth hormone concentrations, SWS proportion, SWA, and high-band aperiodic activity, but negatively with cortisol concentrations and REM sleep proportion measured 40–100 min before. Older participants show higher plasma levels of Aβ1-40, but not Aβ1-42, compared to the younger ones.

Specifically, we found that a single night of normal sleep is accompanied by a 13-15% decrease in Aβ1-40 and Aβ1-42 plasma levels, in line with the reports on an approximate 10% reduction in Aβ, both in blood plasma^16,17,18^ as well as in CSF^3^. Furthermore, we found that plasma levels of Aβ1-40 and Aβ1-42 can be predicted by the amount of SWS (assessed by scoring) and SWA (assessed by EEG power) that was observed around one hour prior to a blood measurement. This is in line with previous observations that total sleep disruption, selective disruption of SWS, or sleep fragmentation (e.g., repetitive sleep interruptions such as in physicians on-call) reduce or abolish the overnight Aβ decrease^2,3,18^. Interestingly, several nights of partial sleep deprivation (4 hours of sleep per night), where REM and total sleep were reduced, but SWS was not, did not change plasma or CSF Aβ levels^19^, indicating that the effect is specific for SWS. Following this literature, we suggest that a positive association between SWS and subsequent peripheral Aβ levels reflects SWS’s contribution to cerebral Aβ clearance (Fig.4).

**Figure 4.**
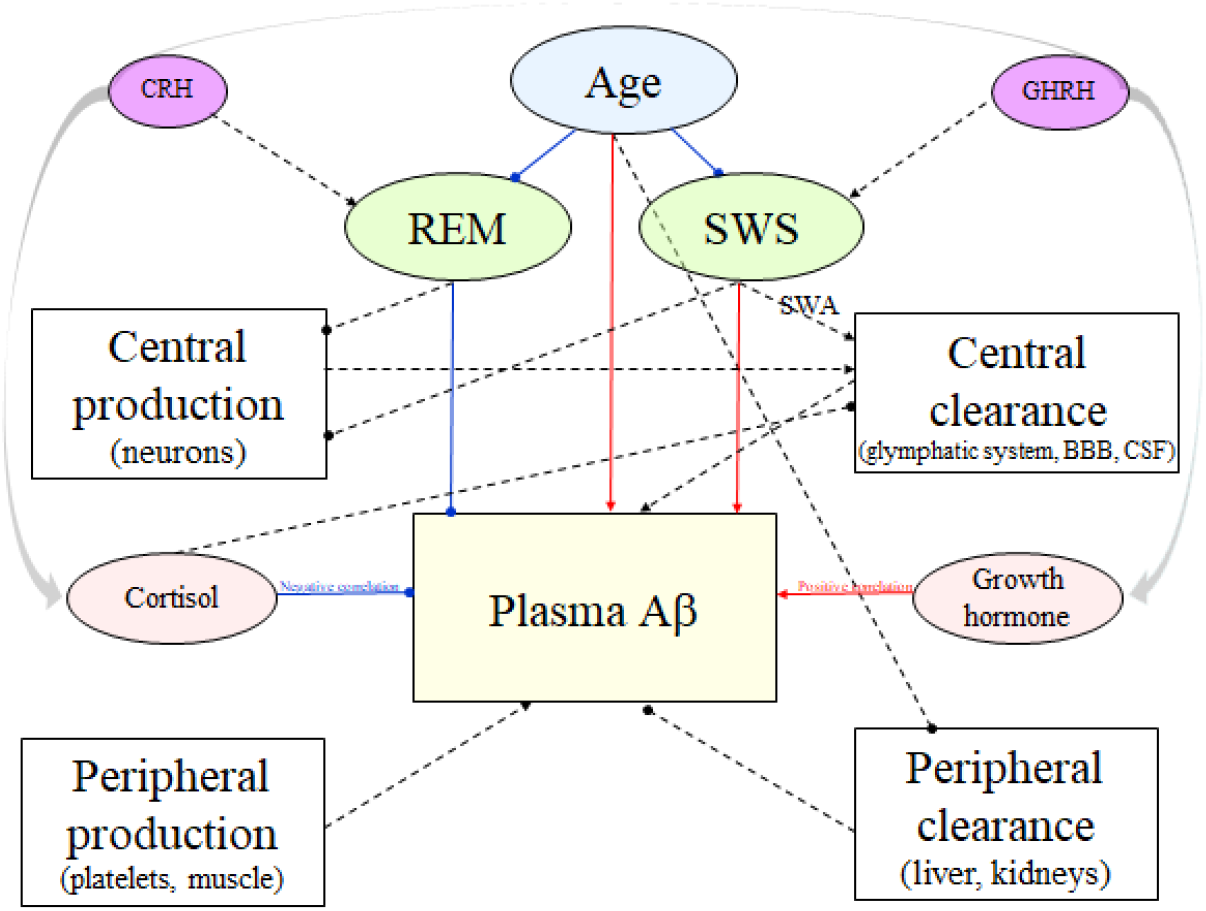
Summary and limitations. SWS contributes to the central clearance of cerebral Aβ (possibly through SWA), leading to higher subsequent plasma Aβ levels. REM sleep is associated with lower subsequent plasma Aβ levels through an unknown mechanism. Healthy aging is associated with decreased SWS and, surprisingly, higher plasma levels of Aβ1-40 (but not Aβ1-42) possibly through inefficient peripheral clearance. It should be kept in mind that Aβ is also synthesized in the periphery (e.g., platelets, muscle, blood vessels) and cleared by the liver. Cortisol negatively correlates with Aβ, whereas growth hormone shows a positive association with Aβ. Previously, it has been shown that GHRH administration increases SWS and growth hormone, whereas CRH elevates cortisol and REM. SWS – slow-wave sleep, SWA-slow-wave activity, REM – rapid eye movements sleep, GHRH – growth hormone-releasing hormone, CRH – corticotrophin-releasing hormone, BBB – blood-brain barrier, CSF – cerebrospinal fluid, red solid arrows – positive association observed in the current study, blue solid arrows – negative association observed in the current study, green lines – non-significant relationships, dashed arrows – associations reported/suggested in the literature.

Following a well-known aging-related decrease in SWS (Fig.S2), one could hypothesize that older adults present reduced brain clearance expressed as lower plasma Aβ. We, however, found similar Aβ1-42 and higher Aβ1-40 in older compared to younger participants (Fig.S3). This is only partially in line with the study by Huang et al.^17^ who found that both Aβ1-40 and Aβ1-42 plasma concentrations are higher in older than in young participants. Nevertheless, in Huang et al.^17^, the correlation between age and Aβ1-40 was considerably higher than between age and Aβ1-42 (r=0.7 vs. r=0.4). It is also worth mentioning that the sample size in Huang et al.^17^ was relatively small compared to our study and comprised only young and old, but not middle-aged participants. Notably, Aβ1-42 is the principal Aβ peptide in Alzheimer’s plaques, whereas Aβ1-40 is a predominant (∼90%) form of Aβ peptide in the brain, CSF, and plasma. Aβ1-40 is not as pathogenic as Aβ1-42 and may even have protective effects against Aβ plaque formation^26^.

A possible explanation for the positive association between age and plasma Aβ1-40 relates to the fact that brain-derived Aβ is also cleared by the liver and kidneys (Fig.4). Following the broadly documented age-related reduction of the renal function^20^, we hypothesize that older persons might have less efficient peripheral Aβ clearance expressed as higher baseline levels of plasma Aβ1-40 (Fig.S3).

Peripheral clearance is very important: according to the peripheral sink hypothesis, cerebral and peripheral soluble Aβ are in equilibrium such that peripheral Aβ clearance induces Aβ removal from the brain into blood^21^. This leads us to the hypothesis that if an age-related decrease in this efflux exists and persists, it can contribute to the cerebral amyloid accumulation and, subsequently, plaque formation and AD development.

The fact that Aβ brain homeostasis is maintained by the brain-periphery equilibrium taken together with the fact that REM sleep typically follows NREM sleep can explain the negative association between REM sleep and subsequent plasma Aβ levels observed here. Presumably, enhanced clearance during SWS results in Aβ accumulation in the blood and thus slows down or even inhibits Aβ efflux from the brain to the periphery during the following sleep stages (Fig.4).

Besides the timing of REM sleep, the negative association between REM and Aβ may be explained by the neural activity during REM. At first glance, this statement is at odds with the very definition of REM sleep as a stage with wake-like desynchronized neural activity, since in such a context, REM sleep should be accompanied by high production of metabolic waste, and a subsequent increase in Aβ plasma levels. The observed relationship, however, suggests that REM sleep may be accompanied by less production, less clearance of waste products, or some combination of both. A possible explanation for this discrepancy might come from the research line studying aperiodic neural activity, i.e., non-oscillatory activity with no defining temporal scale (scale-free) ^14,15^. In both intracranial and scalp human EEG studies, REM and SWS show steeper slopes (faster spectral decay) of the aperiodic power component in the 30–50Hz frequency band as compared to wakefulness^15^. These findings were replicated by our data as well (Fig.S4). Moreover, we found that steeper slopes of high-band – but not low-band – aperiodic activity (a marker of REM sleep) are associated with lower subsequent levels of plasma Aβ. This is in line with the study in a mouse model of Alzheimer’s disease, which has shown that optogenetically inducing gamma (40Hz), but not other frequencies, reduces Aβ peptide levels in the hippocampus^22^.

Recently, it has been suggested that aperiodic slopes reflect the ratio between excitatory and inhibitory neural currents^14,15^. Therefore, steeper aperiodic slopes observed during REM sleep reflect the shift of this ratio in favor of inhibition. A recent study employing magnetic resonance spectroscopy has demonstrated a learning-specific shift towards inhibition during REM sleep^23^. In line with these findings, calcium imaging in rodents has revealed an overall decrease in cortical firing during REM and SWS^24^. Specifically, reduced firing rates during REM sleep are accompanied by a selectively increased activity of inhibitory parvalbumin interneurons. These interneurons mediate reduced cortical activity and increased inhibition of pyramidal neurons^24^, the major source of the EEG signal. Based on these results, Niethard et al. suggest that at the cellular level, REM sleep is a period of prevalent suppression of neural activity where a subset of neurons with high activity during the wake period is activated^24^.

The studies mentioned above contrast with the general view that REM sleep is an active phase of sleep and rather show that REM sleep could be associated with the highest level of cortical inhibition^15,24^. Such a view offers a likely mechanistic explanation for our findings as cortical inhibition should lead to decreased synaptic activity and, consequently, decreased cerebral waste production followed by a lower peripheral Aβ level.

Recently, an fMRI study in naturally sleeping pigeons has shown that REM sleep is associated with a decrease in ventricular CSF flow compared to non-REM sleep, concluding that brain activation during REM sleep comes at the expense of waste clearance^25^.

The assumption that both REM and SWS are associated with cortical inhibition raises the question of why these sleep stages show divergent associations with the subsequent plasma Aβ levels. One possible explanation relates to the timing of REM sleep discussed above (Fig.4). An alternative explanation considers the relative dominance of the oscillatory vs. non-oscillatory neural activity during SWS and REM, respectively^26^. For example, it has been suggested that during SWS, prominent slow oscillatory neuronal activity leads to oscillations in blood volume, drawing cerebrospinal fluid into and out of the brain^27^. Our findings are in line with this theory as here, Aβ fluctuations were associated specifically with slow-wave activity (0.3–4Hz) and not with the oscillatory activity in other frequency bands or broadband (0.3–35Hz) aperiodic activity.

Possibly, all stages of non-REM sleep are involved in brain clearance (with different efficiency) while during REM sleep, clearance is stopped altogether or reduced massively, potentially even compared to wakefulness. If this scenario is true, null associations between REM sleep and brain clearance (the hypothetical ground truth) can be expressed as a negative correlation (the current observation), reflecting REM-related ceasing of clearance, for example, due to the relative dominance of aperiodic compared to oscillatory activity characteristic of this stage^26^, unfavorable brain-periphery Aβ equilibrium or another unknown mechanism.

In summary, a positive effect of SWS on subsequent Aβ plasma levels can be explained by efficient cerebral clearance possibly via the coupling of slow-wave oscillations with blood and cerebrospinal fluid flow. A negative effect of REM sleep on subsequent Aβ might reflect a complex combination of several factors, such as decreased waste production due to neural inhibition, decreased central clearance due to a relative shift to the dominance of non-oscillatory activity or another unknown mechanism, and non-efficient Aβ brain-periphery efflux due to an increased peripheral sink condition observed as an aftereffect of SWS (Fig.4).

In view of the considerable endocrine activity accompanying sleep, we also explored associations between nocturnal fluctuations of hormones and Aβ. We found that Aβ positively correlated with GH and negatively correlated with cortisol plasma levels measured 0–120min before. The broad range of significant time lags can be explained by the fact that cortisol readily crosses the blood-brain barrier and binds to its receptors in the brain^27^. A small but significant amount of GH also passes the blood-brain barrier and acts on the neuronal GH receptors with a modulatory effect on memory, working capacity, and cognition^28^.

In animal models and cell cultures, glucocorticoids have been shown to mediate enhanced production of cerebral Aβ, reduce degradation, facilitate plaque formation, enhance Aβ- mediated neuronal toxicity, and increase tau accumulation^27^ (Fig.S5). Specifically, glucocorticoid treatment *in vitro* and monkeys reduced the activity of the insulin-degrading enzyme, a candidate protease in the clearance of Aβ in the brain, which is decreased in AD patients^27^. Cortisol likely increases the transcription of the amyloid precursor protein (APP) gene via the glucocorticoid-response binding element, leading to the increased Aβ production observed *in vitro* and *in vivo*. An APP protein increase leads to increased processing of APP to C99, which is consequently cleaved by the γ-secretase to release Aβ^27^.

These findings are highly relevant for clinical research because it is established that early sporadic AD patients show elevated cortisol levels, which along with other environmental and genetic risk factors contribute to increased Aβ production and exacerbate existing AD pathologies^27^. Besides AD, elevated cortisol levels can be caused by administration of the exogenous glucocorticoids (e.g., for inflammation treatment) and a stressful lifestyle (Fig.S5), factors that are very common in modern society.

Alternatively, the observed associations can be explained by the timing of the hormone’s release and not by their action on Aβ clearance per se; for example, via the sleep-regulating role of the corresponding releasing hormones (Fig.4). Thus, it is well-documented that the GH-releasing hormone synchronizes both SWS and GH release. The corticotropin-releasing hormone regulates both REM sleep as well as the release of adrenocorticotropin by the pituitary gland and cortisol by the adrenal gland^8^. Undoubtedly, further studies are needed to clarify the mechanisms of the associations between endocrine activity and brain clearance^29^. At this stage, our findings suggest that one’s hormone profile may serve as a new marker of brain clearance efficiency.

This study has several limitations. First, even though plasma is a more accessible and less invasive source than CSF for estimating Aβ concentrations in circulation, peripheral Aβ is not a direct surrogate for central Aβ. Second, one should keep in mind that Aβ is synthesized not only in the brain but also in the periphery (e.g., platelets, muscle, blood vessels)^30^. Circulating Aβ peptides enter the bloodstream and are partly cleared by the liver and kidneys (Fig.4). In addition, the hydrophobic nature of Aβ makes the peptide bind to numerous binding plasma proteins (e.g., ApoE, albumin, Aβ-specific IgG), which could result in “epitope masking” and other analytical interferences^30^. Of note, we used the fluctuations of Aβ plasma concentration as a proxy for central clearance. However, it should be kept in mind that plasma Aβ levels are also influenced by central and peripheral production and peripheral clearance (Fig.4). Plasma analysis cannot differentiate between these sources. Here, we used univariable models with a single sleep feature/hormone and a fixed time lag (but corrected for age and participant-specific random intercepts) rather than multivariable mixed-effects models with multiple sleep features/hormones at several time lags. Our models serve as a descriptive tool, and to gain first insights into the data. However, separate univariable regressions are not directly comparable with each other as they are not based on the exact same set of data points.

Future studies using advanced statistical analyses and multivariable regressions based on our initial descriptive visualizations might be able to overcome this limitation and thereby allow for comparison across time. In addition, it may be interesting to investigate potential non-linearities in predictor variables. Despite these limitations, our study provides a new view on sleep and hormones’ effects on Aβ clearance, suggesting that SWS and REM sleep have different homeostatic functions due to divergent neural and endocrine activity.

### Conclusion

This study demonstrates that specific sleep and hormonal features can predict nocturnal Aβ fluctuations in a sample of healthy volunteers and suggests that simultaneous polysomnography and all-night blood sampling is a promising research tool for investigating brain clearance. Future studies addressing the topic could provide a deeper understanding of the mechanism involved in the pathogenesis of AD and other neurodegenerative conditions, potentially leading to the development of innovative and evidence-based prevention and intervention techniques.

## Contributors

MD and MMV designed the study. AS provided the sleep data and plasma samples. MP, OS, EL, ST, MS handled samples and data. IK, MU and MMV analyzed plasma samples. YR analyzed the data. YR, MP, FW, LB, DAN, NK, TdW, MS, MD discussed analysis strategies. YR, NK and MD wrote the manuscript. All authors contributed to, reviewed, and approved the final draft of the paper. All authors had full access to all the data in the study and had final responsibility for the decision to submit for publication.

## Data sharing

Deidentified individual-level data is shared in Supplementary Material 5. The participants’ privacy has been protected in accordance with applicable laws and regulations. Any additional information required to reanalyze the data reported in this paper is available from the corresponding author upon request.

## Acknowledgments

This research was funded by the Dutch Research Council (NWO), Die Junge Akademie at the Berlin-Brandenburg Academy of Sciences and Humanities and the German National Academy of Sciences Leopoldina. We thank Euroimmun AG, Lübeck, Germany, and ADx NeuroSciences NV, Gent, Belgium, for providing some of the ELISA kits for free.

## Abbreviations

Aβ: amyloid-beta protein
AD: Alzheimer’s disease
AIC: Akaike information criterion
APP: amyloid precursor protein
CSF: cerebrospinal fluid
REM: rapid eye movement
SWA: slow-wave activity
SWS: slow-wave sleep

## Reference

1. Xie L, Kang H, Xu Q, et al. Sleep drives metabolite clearance from the adult brain. Science. 2013;342(6156):373–377.

2. Ooms S, Overeem S, Besse K, Rikkert MO, Verbeek M, Claassen JAHR. Effect of 1 night of total sleep deprivation on cerebrospinal fluid β-amyloid 42 in healthy middle-aged men: a randomized clinical trial. JAMA neurology. 2014;71(8):971–977.

3. Ju YES, Ooms SJ, Sutphen C, et al. Slow wave sleep disruption increases cerebrospinal fluid amyloid-β levels. Brain. 2017;140(8):2104–2111.

4. Varga AW, Wohlleber ME, Giménez S, et al. Reduced slow-wave sleep is associated with high cerebrospinal fluid Aβ42 levels in cognitively normal elderly. Sleep. 2016;39(11):2041–2048.

5. Spira AP, Gamaldo AA, An Y, et al. Self-reported sleep and β-amyloid deposition in community-dwelling older adults. JAMA neurology. 2013;70(12):1537–1543.

6. Ju YES, Lucey BP, Holtzman DM. Sleep and Alzheimer disease pathology—a bidirectional relationship. Nature reviews Neurology. 2014;10(2):115–119.

7. Ouanes S, Popp J. High Cortisol and the Risk of Dementia and Alzheimer’s Disease: A Review of the Literature. Frontiers in Aging Neuroscience. 2019;11. https://www.frontiersin.org/article/10.3389/fnagi.2019.00043

8. Steiger A, Dresler M, Kluge M, Schüssler P. Pathology of sleep, hormones and depression. Pharmacopsychiatry. 2013;46(S01):S30–S35.

9. Lerche S, Brockmann K, Pilotto A, et al. Prospective longitudinal course of cognition in older subjects with mild parkinsonian signs. Alzheimer’s research & therapy. 2016;8(1):1–8.

10. Wen H, Liu Z. Separating fractal and oscillatory components in the power spectrum of neurophysiological signal. Brain topography. 2016;29(1):13–26.

11. Oostenveld R, Fries P, Maris E, Schoffelen JM. FieldTrip: open source software for advanced analysis of MEG, EEG, and invasive electrophysiological data. Computational intelligence and neuroscience. 2011.

12. Rosenblum Y, Bovy L, Weber FD, Steiger A, Zeising M, Dresler M. Increased aperiodic neural activity during sleep in major depressive disorder. Biological Psychiatry: Global Open Science. 2022.

13. Rosenblum Y, Shiner T, Bregman N, Giladi N, Maidan I, Fahoum F, Mirelman A. Decreased aperiodic neural activity in Parkinson’s disease and dementia with Lewy bodies. Journal of Neurology. 2023 May 3:1–2. https://doi.org/10.1007/s00415-023-11728-9

14. Gao R, Peterson EJ, Voytek B. Inferring synaptic excitation/inhibition balance from field potentials. Neuroimage. 2017;158:70–78.

15. Lendner JD, Helfrich RF, Mander BA, et al. An electrophysiological marker of arousal level in humans. Elife. 2020;9:e55092.

16. Benedict C, Blennow K, Zetterberg H, Cedernaes J. Effects of acute sleep loss on diurnal plasma dynamics of CNS health biomarkers in young men. Neurology. 2020;94(11):e1181–e1189.

17. Huang Y, Potter R, Sigurdson W, et al. β-amyloid dynamics in human plasma. Archives of neurology. 2012;69(12):1591–1597.

18. Grimmer T, Laub T, Hapfelmeier A, et al. The overnight reduction of amyloid β 1-42 plasma levels is diminished by the extent of sleep fragmentation, sAPP-β, and APOE ε4 in psychiatrists on call. Alzheimer’s & Dementia. 2020;16(5):759–769.

19. Olsson M, Ärlig J, Hedner J, Blennow K, Zetterberg H. Sleep deprivation and plasma biomarkers for Alzheimer’s disease. Sleep medicine. 2019;57:92–93.

20. Wang YR, Wang QH, Zhang T, et al. Associations Between Hepatic Functions and Plasma Amyloid-Beta Levels—Implications for the Capacity of Liver in Peripheral Amyloid-Beta Clearance. Molecular Neurobiology. 2017;54(3):2338–2344.

21. Roberts KF, Elbert DL, Kasten TP, et al. Amyloid-β efflux from the central nervous system into the plasma. Annals of Neurology. 2014;76(6):837–844.

22. Iaccarino HF, Singer AC, Martorell AJ, Rudenko A, Gao F, Gillingham TZ, Mathys H, Seo J, Kritskiy O, Abdurrob F, Adaikkan C. Gamma frequency entrainment attenuates amyloid load and modifies microglia. Nature. 2016 Dec;540(7632):230–5.

23. Tamaki M, Wang Z, Barnes-Diana T, et al. Complementary contributions of non-REM and REM sleep to visual learning. Nature neuroscience. 2020;23(9):1150–1156.

24. Niethard N, Hasegawa M, Itokazu T, Oyanedel CN, Born J, Sato TR. Sleep-stage-specific regulation of cortical excitation and inhibition. Current biology. 2016;26(20):2739–2749.

25. Ungurean G, Behroozi M, Boeger L, Helluy X, Libourel PA, Gunturkun O, Rattenborg N. Wide-spread brain activation and reduced CSF flow during avian REM sleep.

26. Helfrich RF, Lendner JD, Knight RT. Aperiodic sleep networks promote memory consolidation. Trends in Cognitive Sciences. Published online 2021.

27. Fultz NE, Bonmassar G, Setsompop K, et al. Coupled electrophysiological, hemodynamic, and cerebrospinal fluid oscillations in human sleep. Science. 2019;366(6465):628–631.

28. Green KN, Billings LM, Roozendaal B, McGaugh JL, LaFerla FM. Glucocorticoids increase amyloid-β and tau pathology in a mouse model of Alzheimer’s disease. Journal of Neuroscience. 2006 Aug 30;26(35):9047–56.

29. Åberg ND, Brywe KG, Isgaard J. Aspects of growth hormone and insulin-like growth factor-I related to neuroprotection, regeneration, and functional plasticity in the adult brain. TheScientificWorldJournal. 2006;6:53–80.

30. Bu XL, Xiang Y, Jin WS, et al. Blood-derived amyloid-β protein induces Alzheimer’s disease pathologies. Molecular psychiatry. 2018;23(9):1948–1956.

