## Supplemental Material for "Divergent associations of slow-wave sleep vs. REM sleep with plasma amyloid-beta"

### Supplementary Materials

Yevgenia Rosenblum [1], Mariana Pereira [1], Oliver Stange [1], Frederik D. Weber [1,2], Leonore Bovy [1], Sofia Tzioridou [1], Elisa Lancini [3,4], David A. Neville [1], Nadja Klein [5], Timo de Wolff [6], Mandy Stritzke [6], Iris Kersten [7], Manfred Uhr [8], Jurgen A.H.R. Claassen[1], Axel Steiger [8], Marcel M. Verbeek [7], Martin Dresler [1]

[1] Radboud University Medical Centre, Donders Institute for Brain, Cognition and Behavior, Nijmegen, Netherlands, [2] Netherlands Institute for Neuroscience, Department of Sleep and Cognition, Amsterdam, Netherlands, [3] German Center for Neurodegenerative Diseases, Otto-von-Guericke University Magdeburg, Magdeburg, Germany, [4] Institute of Cognitive Neurology and Dementia Research, Otto-von-Guericke University Magdeburg, Magdeburg, Germany, [5] Chair of Uncertainty Quantification and Statistical Learning, Research Center for Trustworthy Data Science and Security (UA Ruhr) and Department of Statistics (Technische Universität Dortmund), Germany, [6] Technische Universität Braunschweig, Institut für Analysis und Algebra, Braunschweig, Germany, [7] Radboud University Medical Centre, Departments of Neurology and Laboratory Medicine, Nijmegen, Netherlands, [8] Max Planck Institute of Psychiatry, Munich, Germany.

Corresponding author: Yevgenia Rosenblum, Radboud University Medical Centre, Donders Institute for Brain, Cognition and Behavior, Kapittelweg 29, 6525 EN Nijmegen, the Netherlands.

### Supplementary Figures

**A**

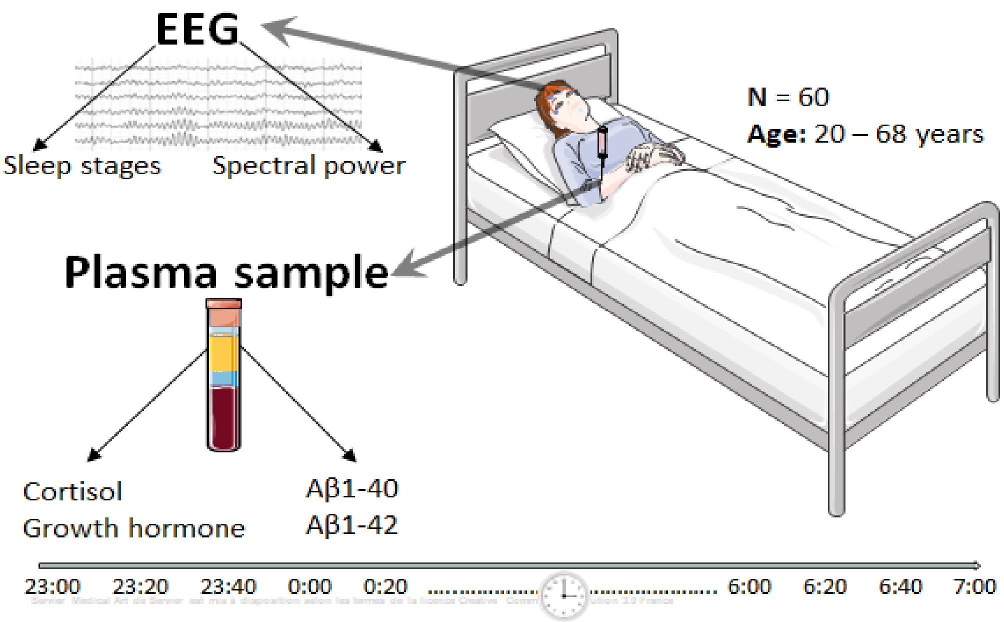


**B**

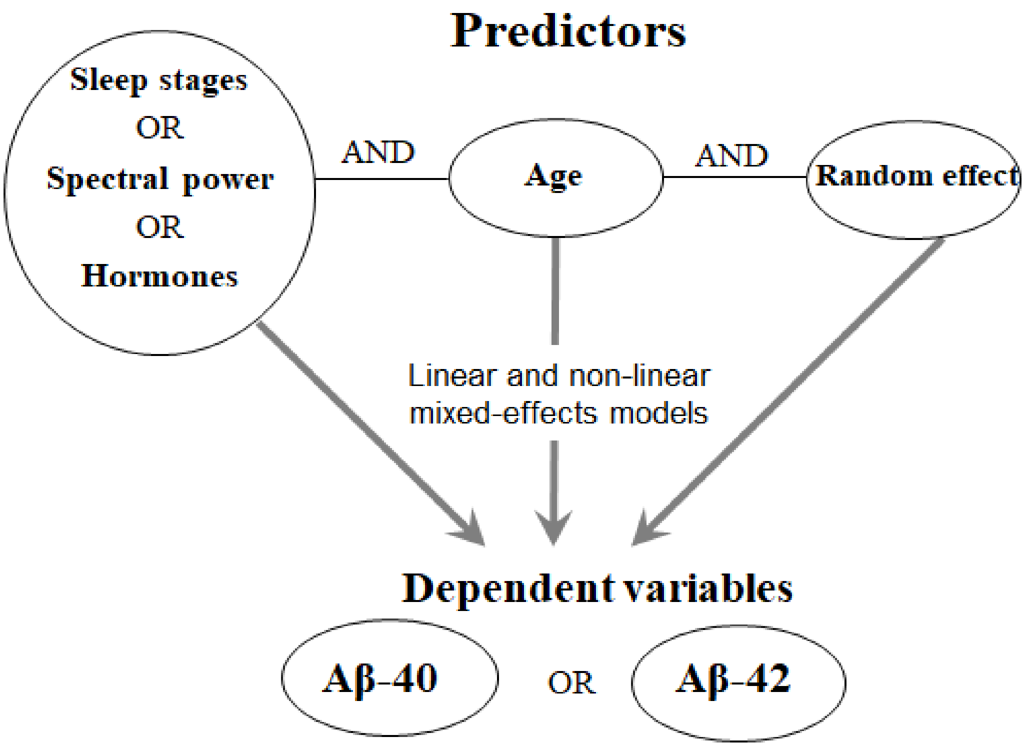

C

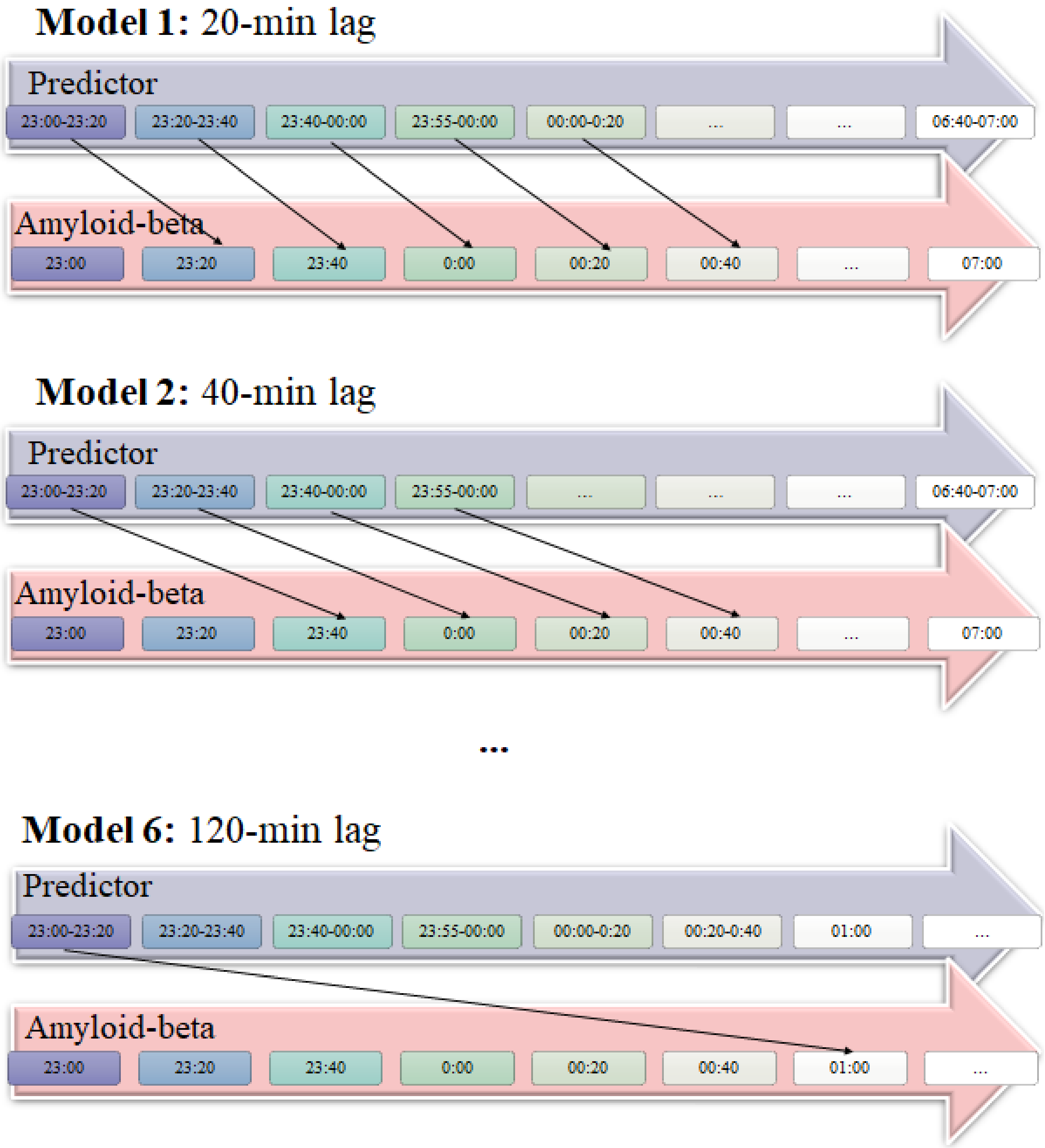


**Supplemental Figure 1. Study design and statistical models. A.** Simultaneous polysomnography and all-night blood sampling were acquired in 60 healthy volunteers aged 20–68 years old. Plasma concentrations of amyloid-beta-40 and amyloid-beta-42, cortisol, and growth hormone were assessed for every 20 minutes of sleep from 23:00 to 7:00. **B.** Predictors and dependent variables used in the mixed models. **C.** For each predictor (sleep/hormone feature), we ran 6 different models corresponding to the time lags between the predictor and Aβ ranging from -120 to -20 min with a 20-min step while controlling for multiple comparisons.

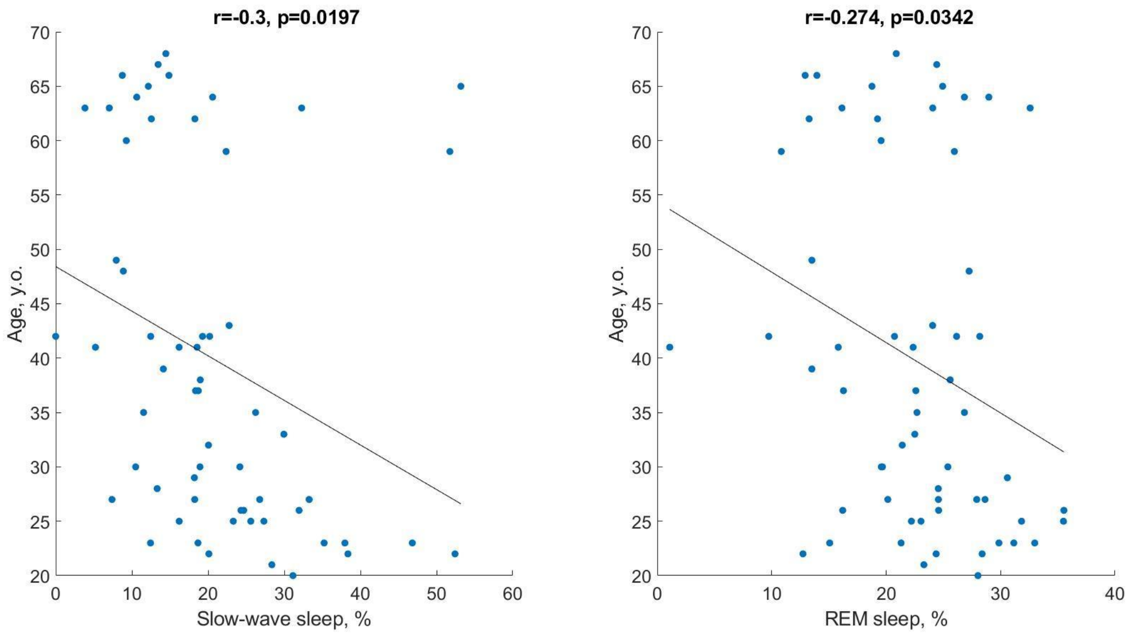


**Supplemental Figure 2. Age, SWS, and REM sleep.** Pearson's correlations between the participants' age on one side and the amount of their SWS and REM sleep on the other side. Both SWS and REM sleep negatively correlate with the participants' age in line with the broad literature, r – Pearson's correlation coefficients, n=60, SWS – slow-wave sleep, REM – rapid eye movement sleep.

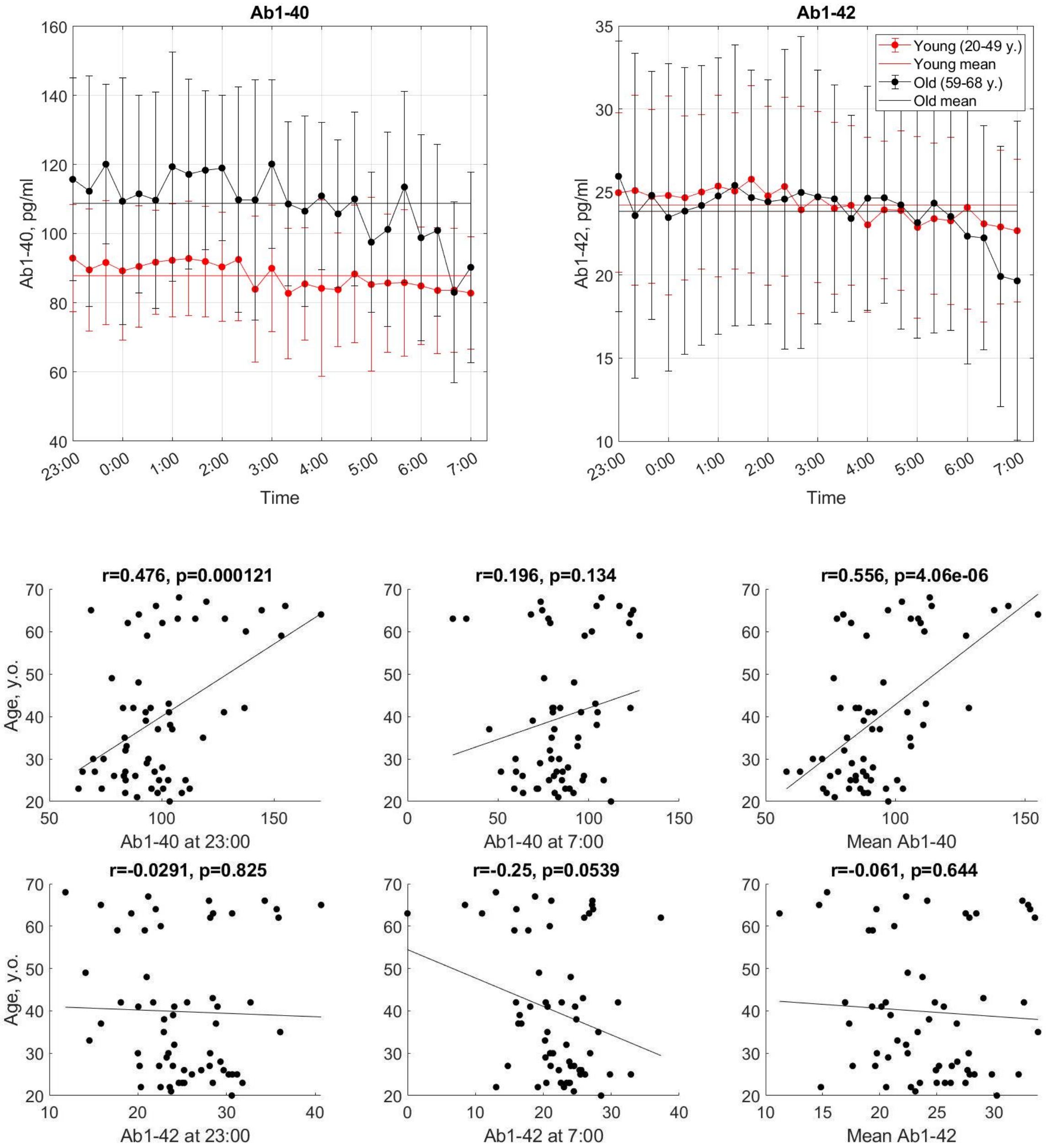


**Supplemental Figure 3. Age and Aβ.** Older (59 – 68 y.o.) participants show higher levels of Aβ1-40 compared to the younger (20 – 49 y.o.) ones. Pearson's correlations between the participants' age on one side and the pre-sleep (measured at 23:00), post-sleep (measured at 7:00) and mean levels of Aβ1-40 and Aβ1-42 on the other side. Exclusively the pre-sleep and mean levels of Aβ1-40 positively correlate with the participants' age, r – Pearson's correlation coefficients, n=60.

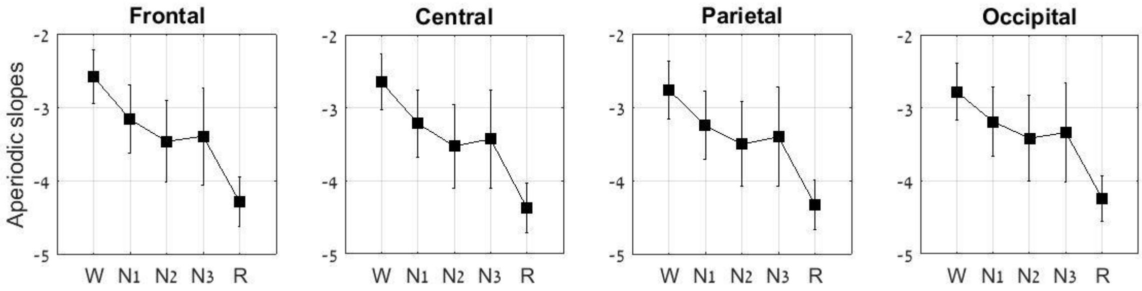


**Supplemental Figure 4. High aperiodic slopes per sleep stage.** This figure visualizes the rationale for using the slopes of the aperiodic spectral power component in the 30–48Hz range as markers of REM sleep. Aperiodic slopes were averaged over each sleep stage as defined by the hypnogram. REM sleep is characterized by more negative (steeper spectral decay) aperiodic slopes compared to both SWS and wakefulness in all topographical areas, n=60, W – wake, N3 – slow-wave sleep, R – rapid eye movement sleep.

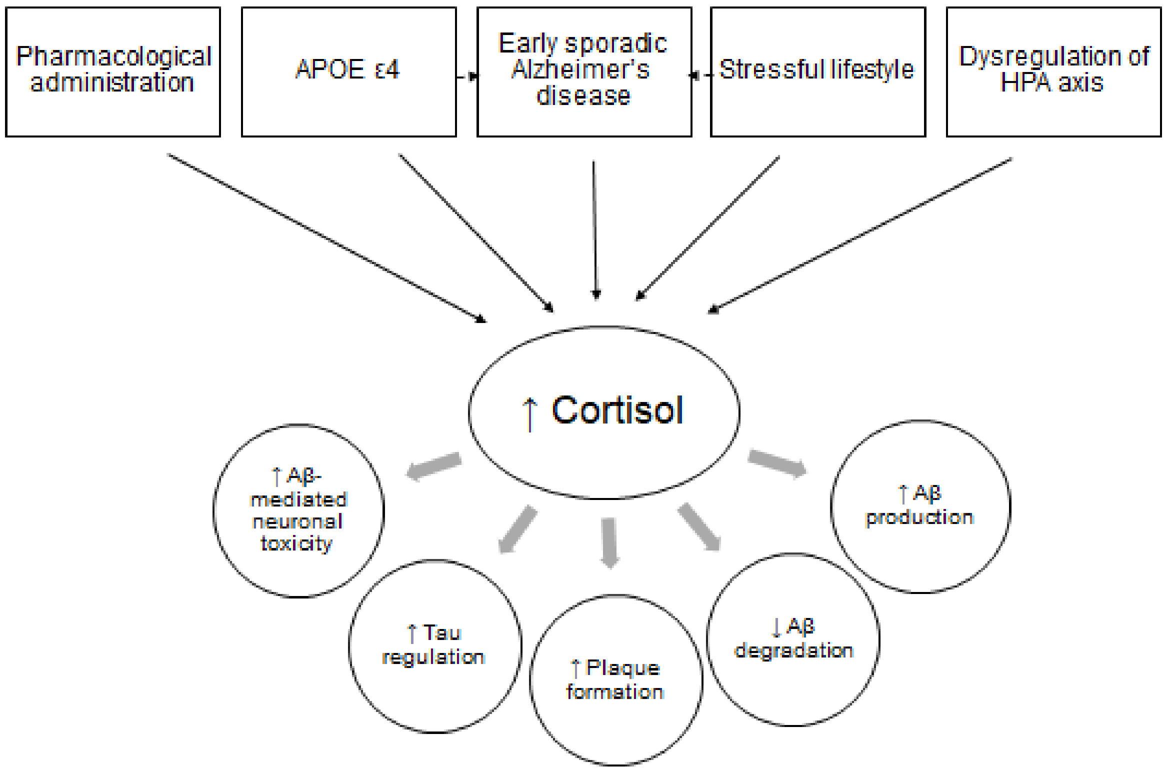


**Supplemental Figure 5. Literature summary on cortisol effects on cerebral Aβ.** Animal models and cell cultures studies revealed that glucocorticoids mediate enhanced production of Aβ, reduce degradation, facilitate plaque formation, enhance Aβ-mediated neuronal toxicity, and increase tau accumulation. In humans, elevated cortisol levels can be caused by early sporadic Alzheimer's disease, the presence of the APOE **ε**4 allele, dysregulation of the HPA axis, administration of exogenous glucocorticoids (e.g., for inflammation treatment), or stressful lifestyle. HPA – hypothalamic–pituitary–adrenal.

### Supplementary Materials

Besides sleep outcome measures reported in the Main text, we also assessed the effects of spectral power over additional topographical areas (S1), atonia (S2), and autonomic functioning (S3) on Aβ clearance. In S4 and S5, we share the analysis R script and the associated dataset, respectively.

#### S1. Topographical analysis

To explore the topography of the effect of the spectral power on Aβ, we calculated the oscillatory and aperiodic power components averaged over 60–80 min prior to a blood measurement (the best quality models as revealed by their AIC) as described in Methods and averaged them over the central (C3, C4), parietal (P3, P4), and occipital (O1, O2) electrodes, separately, in addition to the frontal (F3, F4) electrodes reported in the Main Text.

The topographical analysis revealed that the reported in Results positive correlation between frontal SWA or aperiodic slope in the 30-48Hz band and the subsequent Aβ1-40 and Aβ1-42 plasma levels can be observed over the central, parietal, and occipital electrodes with no prominent topographical effect (S. Table 1).

**Supplemental Table 1: The topography of the main effect of the spectral power on Aβ levels**

| Dependent variable | | Aβ1-40 models | | | | | | Aβ1-42 models | | | | | |
|---|---|---|---|---|---|---|---|---|---|---|---|---|---|
| Predictor | | Slope estimate | SE | t | p | $R^2$ | AIC | Slope estimate | SE | T | p | $R^2$ | AIC |
| SWA | Frontal | 1.370 | 0.352 | 3.891 | <0.001 | 0.641 | 9130 | 0.247 | 0.076 | 3.257 | 0.001 | 0.745 | 5791 |
| | Central | 2.374 | 0.553 | 4.295 | <0.001 | 0.643 | 9126 | 0.447 | 0.118 | 3.775 | 0.000 | 0.746 | 5788 |
| | Parietal | 2.823 | 0.706 | 3.997 | <0.001 | 0.642 | 9129 | 0.489 | 0.151 | 3.231 | 0.001 | 0.745 | 5792 |
| | Occipital | 1.619 | 0.642 | 2.524 | 0.012 | 0.638 | 9138 | 0.282 | 0.138 | 2.051 | 0.040 | 0.744 | 5798 |
| Aperiodic activity | Frontal | 2.385 | 0.748 | 3.187 | 0.001 | 0.640 | 9135 | 0.481 | 0.161 | 2.995 | 0.003 | 0.745 | 5793 |
| | Central | 2.286 | 0.738 | 3.096 | 0.002 | 0.640 | 9135 | 0.446 | 0.159 | 2.808 | 0.005 | 0.744 | 5794 |
| | Parietal | 2.405 | 0.767 | 3.135 | 0.002 | 0.640 | 9135 | 0.452 | 0.165 | 2.739 | 0.006 | 0.744 | 5794 |
| | Occipital | 2.516 | 0.789 | 3.188 | 0.001 | 0.640 | 9135 | 0.502 | 0.170 | 2.957 | 0.003 | 0.745 | 5793 |

*Each row represents the main effect of a fixed factor from an independent mixed-effects model. The time lag between the predictors and Aβ =-80min. Non-significant p-values or p-values that did not survive the correction for multiple comparisons are shown in grey. No. of observations ~1100, No. of subjects=60, SWA – slow-wave activity, SE – standard errors, AIC – Akaike Information Criterion.*

**S2. EMG analysis**

To evaluate a possible effect of atonia on Aβ clearance, we calculated the oscillatory power component and aperiodic slope of the EMG channel filtered in 52–100Hz (as described in Methods) to account purely for muscle activity with no overlapping frequencies with the EEG. The results are presented in Fig. S6-7. We found a positive effect of the aperiodic slopes on Aβ1-40 levels measured at 80–100min thereafter (observed p-value=0.0224 > corrected p-value=0.0083) and a positive effect of the oscillatory component on the Aβ1-40 levels measured at 60–80min thereafter (observed p-value=0.0084 > corrected p-value=0.0083). However, these effects did not pass the correction for multiple comparisons.

We found no effect of the oscillatory power on subsequent Aβ1-42 levels. The aperiodic slopes had a positive effect on Aβ1-42 levels measured at 80–100min (observed p-value=0.0058 < corrected p-value=0.0083) and 100–120min thereafter (observed p-value=0.040 > corrected p-value=0.0166). The latter effect has not survived the correction for multiple comparisons.

The findings show that higher muscle activity to some degree is associated with higher subsequent Aβ levels. A possible interpretation is that muscle activity facilitates peripheral blood flow to the organs of peripheral clearance, namely, the kidneys and liver (Fig.4).

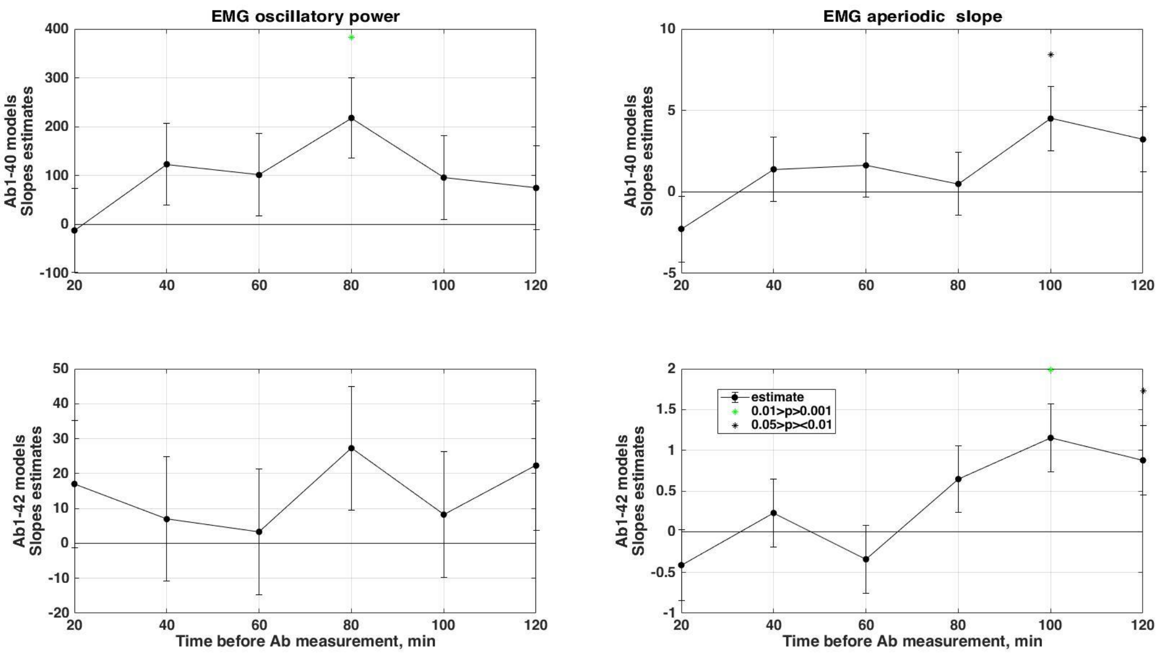

**Supplemental Figure 6. Slope estimates of EMG power.** EMG power components were entered as predictors of Aβ1-40 and Aβ1-42 into 6 different linear mixed-effects models using time lags ranging from -20 to -120 min between the predictors and Aβ. The slope estimate of the fixed effect of each predictor and its SE are presented. The asterisks mark the slope estimates for which the null hypothesis (stating that a predictor does not significantly affect the response (Aβ)) should be rejected. The models that survived the correction for multiple comparisons are listed in the Main Text. Higher oscillatory EMG power predicts higher subsequent Aβ1-40 levels (significantly positive slope) measured 60–80min thereafter. Higher aperiodic slopes predict higher subsequent Aβ1-40 and Aβ1-42 levels measured 80–100min thereafter. Black asterisks correspond to 0.05>p>0.01, green – to 0.01>p>0.001.

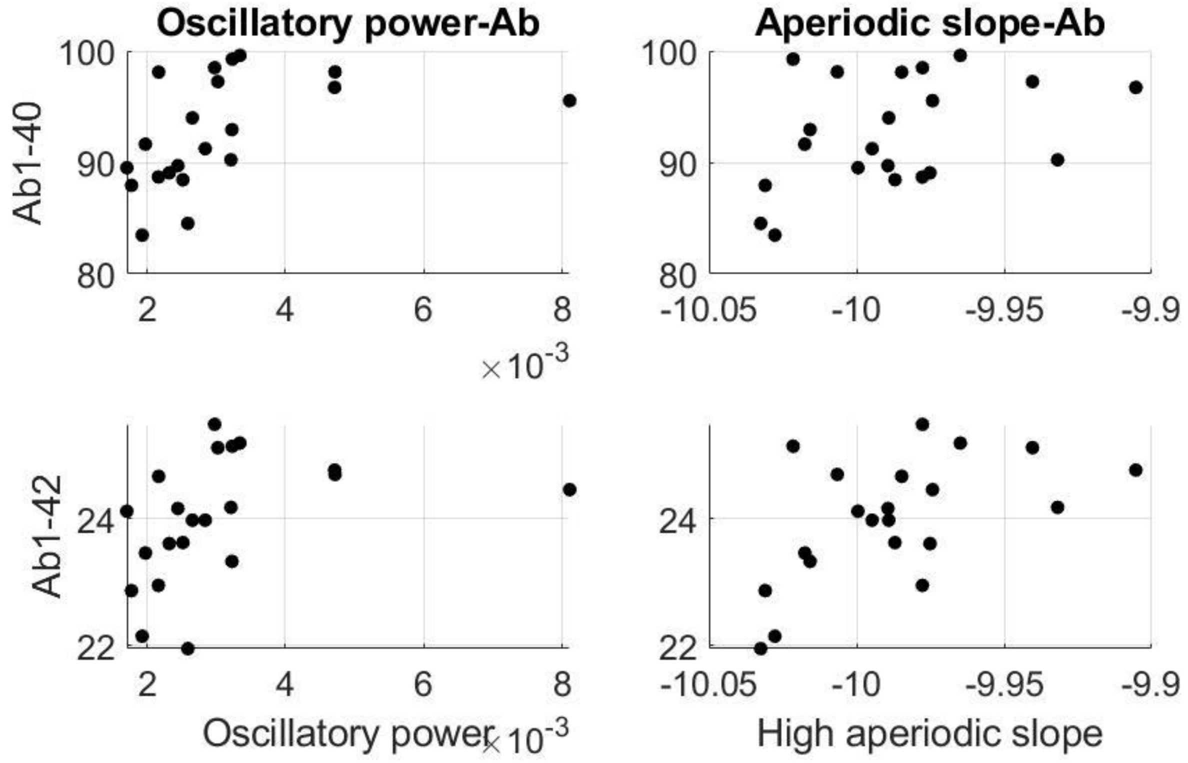


**Supplemental Figure 7. Aβ and EMG.** The relationships between the group-level averaged oscillatory power component (left) or aperiodic slope (right) filtered in 52–100Hz and plasma Aβ levels measured 80 min thereafter.

### S3. ECG analysis

A recent study has suggested that cerebral clearance may depend on the autonomic regulation of CSF pulsations without requiring slow-wave oscillations or even sleep (Piccioni et al., 2022). To evaluate the effect of autonomic functioning on Aβ clearance, here, we calculated the heart rate and heart rate variability (changes in heart rate over time) in the time domain using the *ft_artifact_zvalue* function while filtering the ECG channel of polysomnography in the 20–45Hz range. Specifically, we calculated the average and standard deviation of NN intervals for each 20-min period of sleep, corresponding to each blood measurement. We also calculated spectral power of the ECG channel in the frequency domain as described in the Methods, filtering the signal in the 0.3–0Hz band according to the AASM recommendations.

The results are presented in Fig. S8–9. We found a positive effect of the heart rate on the Aβ1-42 levels measured at 20–40min and 80–120min thereafter (p-values=0.019–0.023). However, only the effect observed at 20–40min passed the correction for multiple comparisons (observed p-value=0.023 < corrected p-value=0.025). There was no effect of spectral power of the ECG or heart rate variability on subsequent Aβ levels.

The findings indicate that a higher heart rate is associated with higher subsequent Aβ1-42 levels. A higher heart rate may reflect autonomic arousal or autonomic instability, which, in turn, may result in CSF pulsations that mediate clearance (Piccioni et al., 2022).

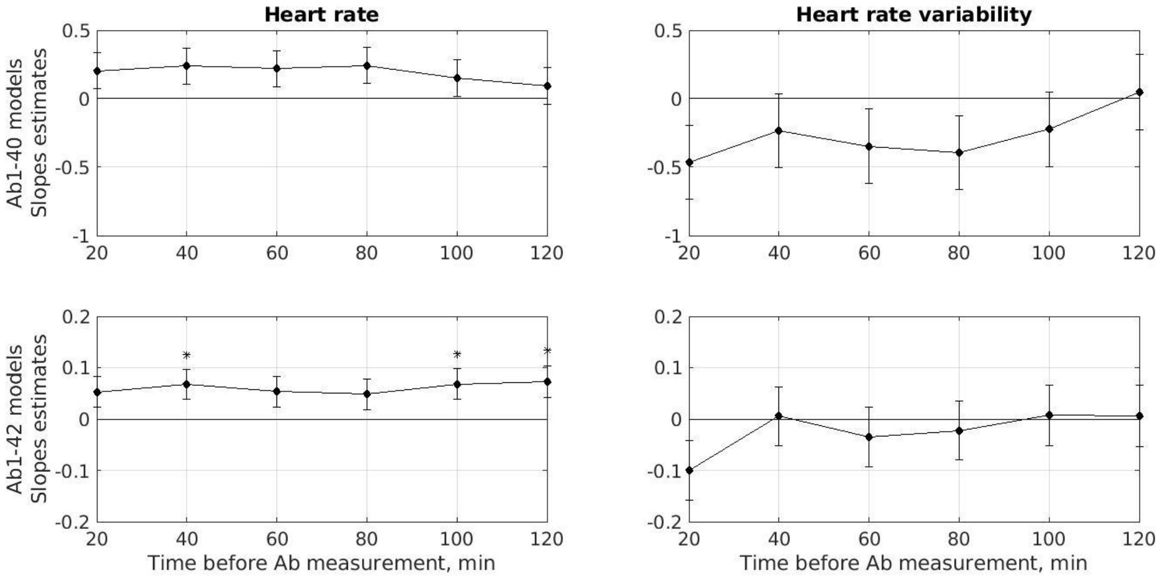


**Supplemental Figure 8. Slope estimates of the heart rate.** The heart rate and heart rate variability were entered as Aβ predictors into 6 different linear mixed-effects models using time lags ranging from -20 to -120 min between the predictors and Aβ. The slope estimate of the fixed effect of each predictor and its SE are presented. The asterisks mark the slope estimates for which the null hypothesis (stating that a predictor does not significantly affect the response (Aβ)) should be rejected. The models that survived the correction for multiple comparisons are listed in the Main Text. Higher heart rate predicts higher subsequent Aβ1-42 levels (significantly positive slopes) measured 20–40min and 80–120min thereafter, whereas heart rate variability has no effect (close to zero slopes) on the subsequent Aβ levels. Black asterisks correspond to 0.05>p>0.01.

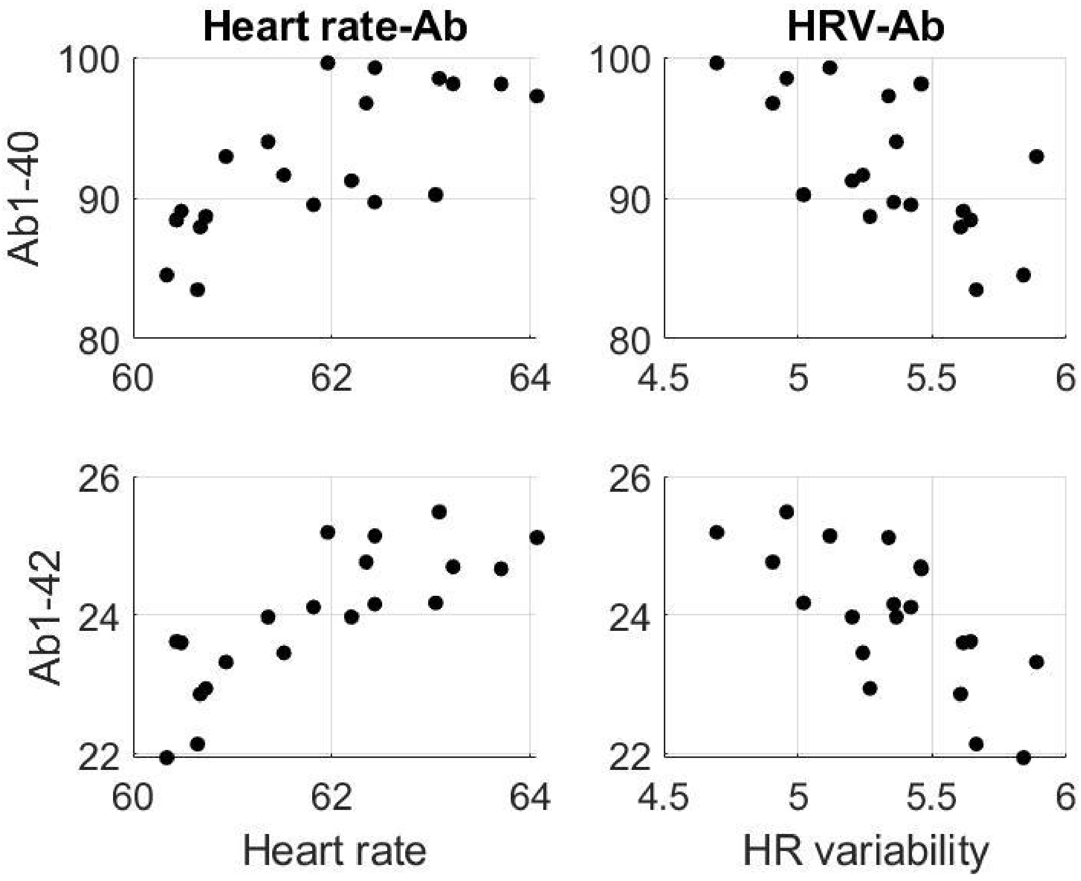


**Supplemental Figure 9. Aβ and ECG.** The relationships between the group-level averaged heart rate (left) or heart rate variability (right) and plasma Aβ levels measured 80–100 min thereafter. HR – heart rate, HRV – heart rate variability.

### S4. R script

Below we present the R script to run linear mixed-effects models with sleep features/hormones, age, and the random effect ("re") of the participants as predictors of the subsequent levels of plasma Aβ-40 or Aβ-42.

```
# Script beginning
rm(list=ls(all=TRUE))

# Open the needed libraries
library(mgcv)
library(eeptools)
```

**# Read the data**

```
data_path = ".../Ab_dataset.csv" # add your path
data = read.csv2(data_path, header=TRUE, sep=",")
```

**# Create lagged columns of Ab40 and Ab42**

```
# use the lag=4, which corresponds to the -80min lag between the predictors and Ab
data <- lag_data(data,group="Subject", time="Time", values=c("Ab42","Ab40"), periods=-4)

run_ab_models <- function(ab_input){
```

**# Create data frame**

```
sleep_ab = data.frame(
        "Subject" = as.factor(data$Subject),
        "Age"=as.numeric(data$Age),
        "Ab" = as.numeric(get(ab_input, data))
  )
```

**# Linear mixed-effects model (lmer)**

```
# age
model_lmer= gam(Ab ~ Age + s(Subject, bs='re'), data= sleep_ab)
table=summary(model_lmer)
table_lmer=matrix(c(table$p.coeff["Age"],       table$se["Age"],       table$p.t["Age"],       table$p.pv["Age"],
table$r.sq,model_lmer$aic), nrow=1, ncol=6, byrow=TRUE)


# We are interested to perform separate models for the following predictors: N1, N2, SWS, REM, SWA, aperiodic slope,
# cortisol, GH (columns 4:11), therefore we will loop through these columns
col_names = c("N1", "N2", "N3", "REM", "SWA", "Aper", "Cortisol", "GH")

for (sleep_info in col_names) {
        sleep_ab = data.frame(
        "Subject" = as.factor(data$Subject),
        "Age"=as.numeric(data$Age),
        "Sleep" = as.numeric(get(sleep_info, data)),
        "Ab" = as.numeric(get(ab_input, data))
        )
```

**# Linear mixed-effects model (lmer)**

```
model_lmer= gam(Ab ~ Sleep + Age + s(Subject, bs='re'), data= sleep_ab)
table=summary(model_lmer)
table_lmer=rbind(table_lmer,matrix(c(table$p.coeff["Sleep"],table$se["Sleep"],table$p.t["Sleep"],table$p.pv["Sleep"],tabl
e$r.sq,model_lmer$aic),nrow=1,ncol=6,byrow=TRUE))
```

```
# Summarizing the results
# create a table summarizing the linear models
colnames(table_lmer) <- c('Slope estimate','SE','t','p','R2','AIC')
rownames(table_lmer) <- c('Age','N1', 'N2', 'N3', 'REM', 'SWA', 'Aper', 'Cortisol','GH')
table_lmer <- as.table(table_lmer)

# save the tables as *.txt file
write.table(table_lmer, paste0(strsplit(ab_input, "[.]")[[1]][1], '_table_lmer.txt'),
            append = FALSE, sep = "\t", dec = ".", row.names = TRUE, col.names = TRUE)
}

# Run the function for each Ab type
run_ab_models("Ab42.lead4")
run_ab_models("Ab40.lead4")
# The end of the script
```

### Reference


Picchioni D, Özbay PS, Mandelkow H, de Zwart JA, Wang Y, van Gelderen P, Duyn JH. Autonomic arousals contribute to brain fluid pulsations during sleep. *NeuroImage.* 2022 Jan 10:118888. https://doi.org/10.1016/j.neuroimage.2022.118888
